# Mechanotaxis directs *Pseudomonas aeruginosa* twitching motility

**DOI:** 10.1101/2021.01.26.428277

**Authors:** Marco J. Kühn, Lorenzo Talà, Yuki Inclan, Ramiro Patino, Xavier Pierrat, Iscia Vos, Zainebe Al-Mayyah, Henriette MacMillan, Jose Negrete, Joanne N. Engel, Alexandre Persat

**Author notes:** Corresponding authors, Alexandre Persat, Joanne Engel. These authors contributed equally (Marco J. Kühn, Lorenzo Talà).

## Abstract

The opportunistic pathogen *Pseudomonas aeruginosa* explores surfaces using twitching motility powered by retractile extracellular filaments called type IV pili. Single cells twitch by successive pili extension, attachment and retraction. However, whether and how single cells control twitching migration remains unclear. We discovered that *P. aeruginosa* actively directs twitching in the direction of mechanical input from type IV pili, in a process we call mechanotaxis. The Chp chemotaxis-like system controls the balance of forward and reverse twitching migration of single cells in response to the mechanical signal. On surfaces, Chp senses type IV pili attachment at one pole thereby sensing a spatially-resolved signal. As a result, the Chp response regulators PilG and PilH control the polarization of the extension motor PilB. PilG stimulates polarization favoring forward migration, while PilH inhibits polarization inducing reversal. Subcellular segregation of PilG and PilH efficiently orchestrates their antagonistic functions, ultimately enabling rapid reversals upon perturbations. This distinct localization of response regulators establishes a signaling landscape known as local-excitation, global-inhibition in higher order organisms, identifying a conserved strategy to transduce spatially-resolved signals. Our discovery finally resolves the function of the Chp system and expands our view of the signals regulating motility.

## Introduction

Single-cell organisms have evolved motility machineries to explore their environments. For example, bacteria utilize swimming motility to propel themselves through fluids. In their natural environments, bacteria are however most commonly found associated to surfaces^1^. Many species use surface-specific motility systems such as twitching, gliding, and swarming to migrate on solid substrates^2^. However, we still know very little about how cells regulate and control surface motility. In particular, the role of mechanical signals in regulating the motility of single cells remains vastly underexplored in bacteria, as well as in higher order microorganisms^3^.

To migrate towards nutrients and light or away from predators and toxins, cells actively steer motility in response to environmental signals. For example, chemotactic systems mediate motility towards specific molecular ligands^4,5^. Bacteria have a remarkably diverse set of chemotaxis systems. The canonical Che system, which has been extensively studied as a regulator of *Escherichia coli* swimming, is widely conserved among motile species including non-swimming ones^6^. However, the signal inputs and the motility outputs of bacterial chemotaxis-like systems remain unidentified in many species^7^.

*Pseudomonas aeruginosa* is a major opportunistic pathogen well-adapted to growth on surfaces. *P. aeruginosa* colonizes and explores abiotic and host surfaces using twitching motility, which is powered by retractile extracellular filaments called type IV pili (T4P)^8^. During twitching, single cells pull themselves by successive rounds of T4P extension, attachment and retraction^8,9^. T4P extend and retract from the cell surface by respective polymerization and depolymerization of the pilin subunit PilA at the poles^8,9^. While an understanding of the assembly mechanisms of individual filaments is beginning to emerge, we still don’t know whether and how cells coordinate multiple T4P at their surface to power migration over large distances.

Several chemical compounds bias collective or single cell twitching migration^10–12^. It however remains unclear whether they passively bias twitching displacements or actively guide motility. Genetic studies suggest that a chemotaxis-like system called Chp regulates twitching^13^. Beyond playing a role in the transcription of T4P genes, the mechanism by which Chp regulates motility remains unknown^14,15^. In addition, unlike homologs from the well-studied canonical *E. coli* Che system, the Chp methyl accepting chemotaxis protein called PilJ has no clear chemical ligand^15,16^. Also unlike Che which possesses a single response regulator, the Chp system possesses two response regulators, PilG and PilH, whose respective functions remain unresolved^16^.

We previously demonstrated that *P. aeruginosa* upregulates genes coding for virulence factors upon surface contact in a T4P-and Chp-dependent manner^17,14^. There is however no clear evidence that Chp controls any other cellular process than transcription, which is unexpected for a chemotaxis system^13,15,18^. The homology between Chp and Che systems suggests a tactic function for Chp. As a result, we rigorously tested the hypothesis that Chp regulates twitching motility of single cells in response to T4P mechanical input at the timescale of seconds.

## Main

The canonical Che system regulates bacterial swimming by transducing an input chemical signal into a motility response via flagellar rotation control^19^. By analogy, we hypothesized that the chemotaxis-like Chp system regulates the trajectory of single twitching cells^15^. Chp mutants twitch aberrantly in the traditional stab assay (Extended Data Fig. 1ab)^14,18^. These mutants also have altered cyclic AMP (cAMP) levels (Extended Data Fig. 1c)^17^. cAMP regulates the transcription of virulence genes upon surface contact, so that Chp mutants have aberrant T4P numbers (Extended Data Fig. 1d)^17^. To overcome a potential cross-talk, we decoupled the Chp-dependent, short timescale motility control from cAMP-dependent transcription by investigating single cell twitching in constitutively low or high cAMP regimes.

**Figure 1.**
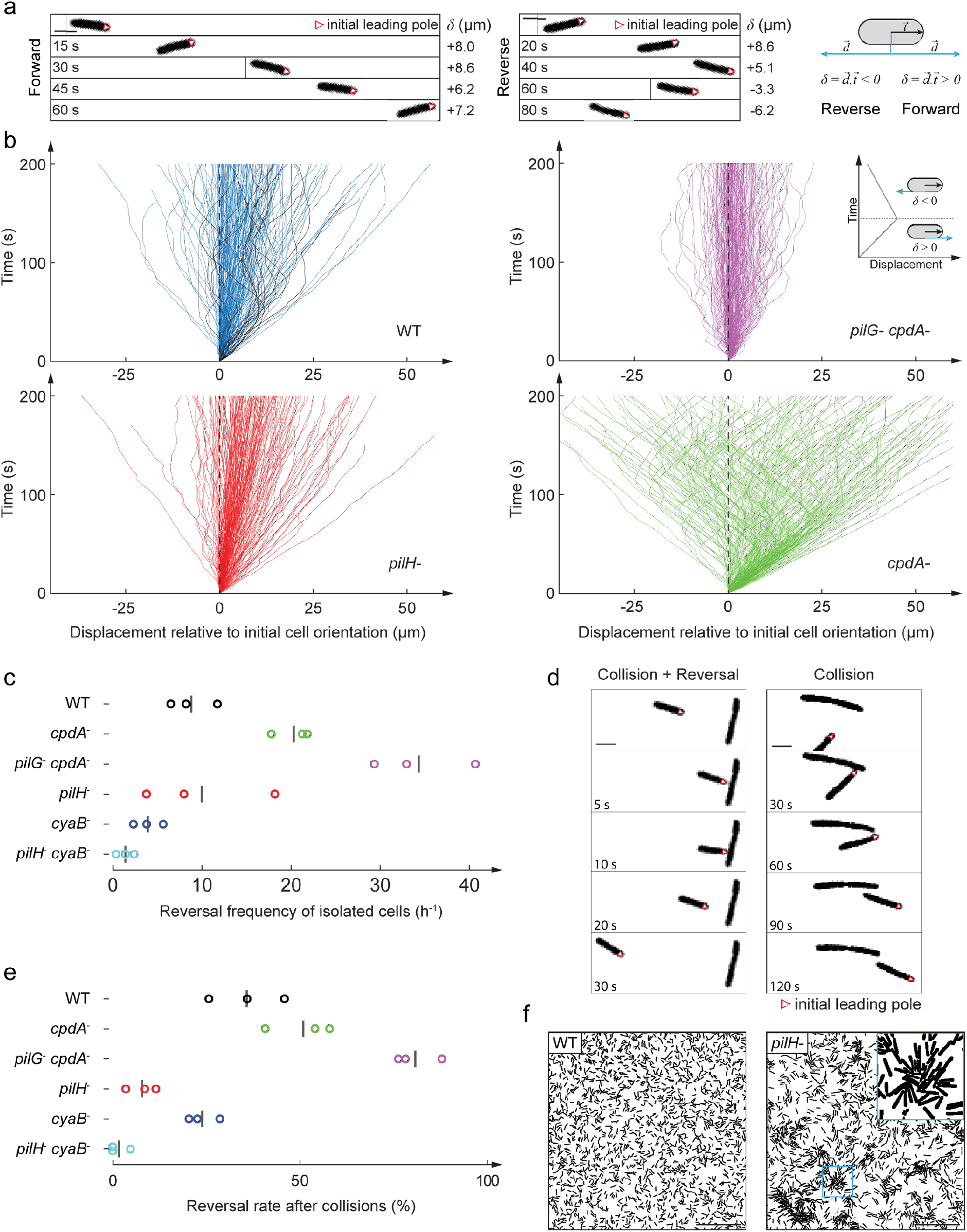
The Chp system regulates the twitching trajectories of individual *P. aeruginosa* cells. (**a**) Phase contrast snapshots of forward and reverse migration. 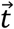 is a unit vector oriented along the cell body in the initial direction of motion. 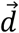 is the unit displacement vector. δis the dot product 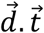, which quantifies displacements relative to the initial direction of motility. Scale bar, 2 μm. (**b**) Graphs of cumulative net displacement as a function of time, highlighting the forward and reverse twitching behavior of Chp mutants. Each curve corresponds to an individual cell trajectory. Tracks of reversing WT cells are highlighted in black. At any given time, a curve oriented toward the top right corresponds to a cell moving forward, while a curve oriented toward top left corresponds to reverse movement (*cf*. inset). *pilG^-^ cpdA^-^* constantly reverses twitching direction while *pilH^-^* cells persistently move forward. (**c**) Quantification of reversal rates in Chp and cAMP mutants. *pilG^-^ cpdA^-^* has highest reversal frequency. *pilH^-^* has a two-fold lower reversal frequency than *cpdA^-^*. Circles correspond to biological replicates, black bars represent their mean. (**d**) Snapshots of WT reversing upon collision with another cell (left). The same sequence for a *pilH^-^* cell, failing to reverse upon collision (right). Scale bar, 2 μm. (**e**) Fraction of cells reversing upon collision with another cell. About half of WT cells reverse after collision, *pilH^-^* almost never reserves after collision, and *pilG^-^ cpdA^-^* almost always reverses. Circles correspond to biological replicates, black bars represent their mean. (**F**) While WT is able to move efficiently at high density, the reduced ability of *pilH^-^* to reverse upon collision leads to cell jamming and clustering. Scale bar, 50 μm. Background strain: PAO1 *fliC^-^*.

We first explored the functions of Chp in directing twitching motility by visualizing individual isolated motile WT (Fig. 1a, Supplementary Video 1), *cpdA^-^*, *pilH^-^*, and *pilG^-^ cpdA^-^* cells, all of which have elevated cAMP levels (Extended Data Fig.1c), at the interface between a glass coverslip and an agarose pad. In all strains, we computed the linear displacements of single cells to visually highlight the balance between persistent forward motion and reversals for single cells (Fig. 1b). We also computed their mean reversal frequency (Fig. 1c). WT and *cpdA^-^* cells mostly move persistently forward and only occasionally reversed twitching direction (Fig. 1bc). *pilG^-^ cpdA^-^* cells reversed so frequently that they appeared to “jiggle”, never really persisting in a single direction of twitching (Supplementary Video 2, Fig. 1bc). They ultimately had very little net migration, consistent with their reduced twitching motility in the stab assay (Extended Data Fig. 1a). By contrast, *pilH^-^* moved very persistently in a single direction and reversed very rarely (Fig. 1bc). Likewise, *pilH^-^ cyaB^-^* with reduced cAMP levels compared to *pilH^-^* had near zero reversal frequency, confirming the Chp-dependent, cAMP-independent control of twitching direction (Fig. 1c).

Upon colliding other cells, WT cells often reversed their twitching direction (Fig. 1d, Supplementary Video 3). *pilG^-^ cpdA^-^* reversed almost always after collision, whereas *pilH^-^* almost never did (Fig. 1ce, Supplementary Video 3 and 4). Because *pilH^-^* cells were unable to reverse, they gradually formed groups by jamming, while WT *P. aeruginosa* were able to spread more evenly (Fig. 1f and Supplementary Video 4). Therefore, Chp provides single *P. aeruginosa* cells with the ability to migrate persistently in one direction and to rapidly change twitching direction. PilG promotes persistent forward motility, driving migration over long distances. PilH enables directional changes particularly useful upon collisions with other bacteria. By controlling reversal rates upon collision, Chp-dependent mechanosensing can enhance the motility of *P. aeruginosa* groups, evoking the control of collective motile behavior in the bacterium *Myxococcus xanthus*^20,21^.

To investigate how PilG and PilH control twitching direction, we focused on the distribution of T4P between the two poles of a cell. We imaged *P. aeruginosa* by interferometric scattering microscopy (iSCAT) to quantify T4P at each cell pole and evaluate their distributions. We found that WT and *cpdA^-^* had T4P distributions close to a random distribution (Extended Data Fig. 2). In contrast, T4P of *pilG^-^ cpdA^-^* were distributed more symmetrically compared to the random distribution and to WT. While *pilH^-^* had too many T4P for a direct comparison with other mutants, we could quantify distributions in the less piliated *pilH^-^ cyaB^-^* background (Extended Data Fig. 1d). The T4P distribution of *pilH^-^ cyaB^-^* was markedly more asymmetric than the random distribution (Extended Data Fig. 2), consistent with its inability to reverse twitching direction. We conclude that the Chp system polarizes T4P to regulate twitching direction. PilG promotes unipolar T4P deployment driving persistent forward migration, while PilH promotes T4P deployment at both poles simultaneously, favoring reversals.

**Figure 2.**
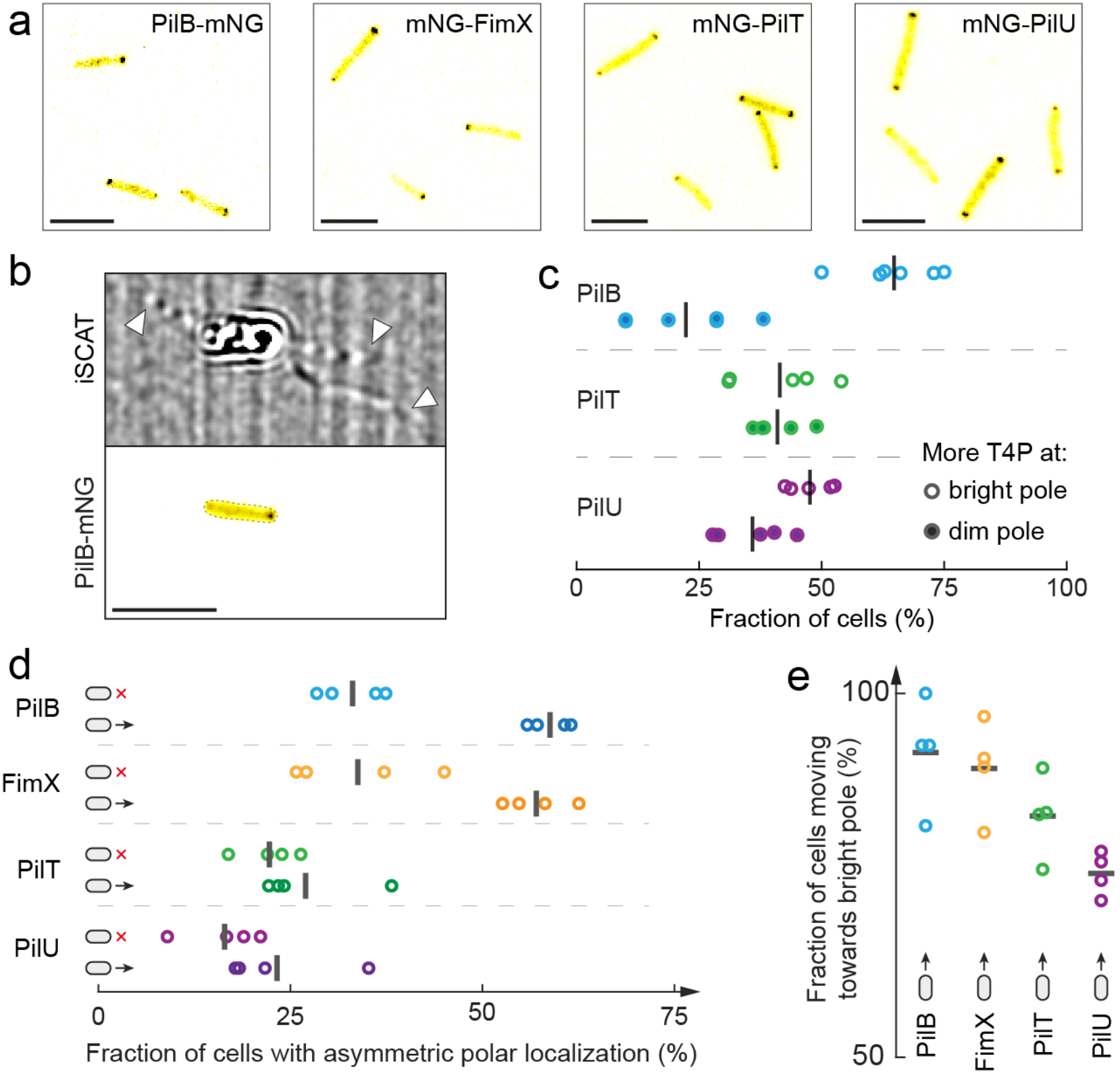
The localization of the extension motor PilB sets the direction of twitching and the polarization of T4P activity. (**a**) Snapshot of chromosomal fluorescent protein fusions to the extension motor PilB, its regulator FimX, and the retraction motors PilT and PilU. Scale bars, 5 μm. (**b**) Simultaneous imaging of PilB-mNG and T4P by correlative iSCAT fluorescence. White arrowheads indicate T4P. Scale bar, 5 μm. (**c**) Fraction of cells with more T4P at bright *vs* dim fluorescent pole. Most cells have more T4P at the bright PilB-mNG pole. We could not distinguish a T4P depletion at the bright retraction motor poles. Each circle is the mean fraction for one biological replicate. Black bars correspond to their mean across replicates. (**d**) Comparison of the symmetry of polar fluorescence between moving and non-moving cells. PilB and FimX signal is more asymmetric in moving cells, which is not the case for PilT and PilU. (**e**) Fraction of cell twitching in the direction of their brightest pole. Circles correspond the fraction of each biological replicate, black bars represent their mean.

T4P extend and retract from the cell surface by respective polymerization and depolymerization of the pilin subunit PilA at the poles. The extension motor PilB assembles PilA monomers to extend T4P, while the retraction motors PilT and PilU disassemble filaments to generate traction forces^8,9^. We reasoned that, for the Chp system to regulate T4P polarization and sets a cell’s twitching direction, it must control either extension or retraction at a given pole. To test this hypothesis, we investigated how the localization of extension and retraction motors regulate the deployment of T4P to direct twitching. First, we simultaneously visualized T4P distribution and motor protein subcellular localization within single cells. To this end, we generated chromosomal mNeonGreen (mNG) fluorescent protein fusions to the extension motor PilB, to its regulator FimX^22^, and to the retraction motors PilT and PilU at their native loci (Fig. 2a). All fusion proteins primarily exhibited bright fluorescent foci at one or both poles consistent with inducible plasmid-borne fusions^23,24^. We leveraged correlative iSCAT-fluorescence microscopy for simultaneous imaging of extracellular filaments and fluorescent reporter fusions (Fig. 2b)^25^. In single cells, we identified the pole with brightest fluorescent signal and the pole with most T4P. We then categorized cells into two groups: cells with more T4P at the bright pole, and cells with less T4P at the bright pole. We found that in more than 60% of cells, the poles with more T4P had the brightest PilB-mNG fluorescent signal (Fig. 2c)^23^. On the other hand, we found no negative correlation between mNG-PilT or mNG-PilU signals and relative numbers of T4P, which would be expected if cells controlled T4P distribution using retraction. As a result, we found that PilB, but neither PilT nor PilU, control the polarized deployment of T4P.

To test whether the control of T4P polarization by PilB ultimately determined *P. aeruginosa* twitching direction, we investigated the dynamic localization motors in single twitching cells (Extended Data Fig. 3, Supplementary Video 5). While mNG-PilT and mNG-PilU fusions were fully functional, PilB-mNG exhibited a partial twitching motility defect (Extended Data Fig. 4a). We therefore systematically validated PilB localization results by visualization of its regulator FimX using mNG-FimX, which was fully functional (Extended Data Fig. 4). We tracked single cells while measuring the subcellular localization of the fusion proteins. We first categorized cells as moving and non-moving. We then measured the proportion of cells that had asymmetric and symmetric protein localizations based on a threshold of fluorescence ratio between poles. We found that PilB-mNG and mNG-FimX localizations were more asymmetric (*i.e*. polarized) in moving cells compared to non-moving cells (Fig. 2d). In addition, both fusion proteins changed localization and polarity during reversals (Extended Data Fig. 5, Supplementary Video 6)^22^. In contrast, the localization of mNG-PilT and mNG-PilU was largely symmetric across the population, without marked symmetry differences between non-moving and moving cells. Since PilB and FimX polarize in moving cells, we computed the correlation between the twitching direction and fusion protein polarization (*i.e*. the localization of their brightest polar spot). We found that more than 90% of cells moved in the direction of the bright PilB and FimX pole (Fig. 2e). Altogether, our data shows that polarized extension and constitutive retraction controls *P. aeruginosa* twitching direction.

**Figure 3.**
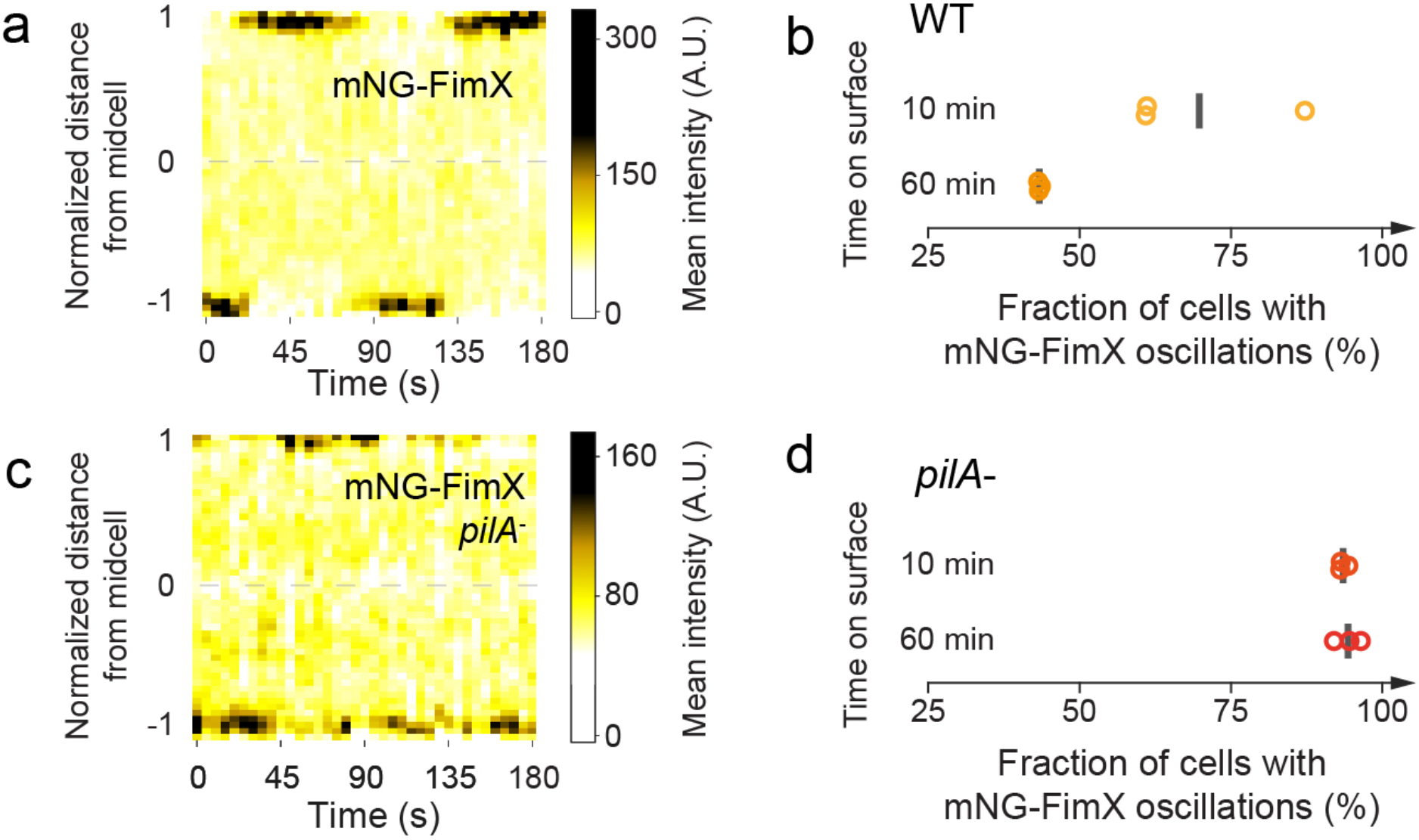
Mechanical input signal from T4P controls the polarization of FimX, the activator of the extension motor PilB. (**a**) Kymograph of mNG-FimX fluorescence in a non-moving cell 10 min after surface contact. The bright fluorescent focus sequentially disappears from one pole to appear at the opposite to establish oscillations. (**b**) Fraction of cells that showed pole to pole oscillations in WT and *pilA^-^*. The proportion of oscillating WT reduces as they remain on the surface, conversely increasing the proportion of stably polarized cells. (**c**) Kymograph of mNG-FimX fluorescence in a *pilA^-^* background 60 min after surface contact. Scale bar, 5 μm. (**d**) Most *pilA^-^* cells maintain oscillatory fluctuations in mNG-FimX polar localization.

T4P mediate a Chp- and cAMP-dependent transcriptional response to surface contact^17^. As a result, we tested whether T4P activity itself regulates PilB polarization. We reasoned that the longer a cell resides on a surface, the more likely it is to experience mechanical stimuli from T4P. We thus compared polarization of cell populations right after contact (10 min) with populations that were associated with the surface for longer times (60 min). We focused on the dynamic localization of mNG-FimX. First, we found that in many cells, polar mNG-FimX foci relocated from pole to pole within a short timeframe after surface contact, as if they were oscillating (Fig. 3a, Supplementary Video 7). These were reminiscent of oscillations in the twitching and gliding regulators observed in *M. xanthus*^21,26,27^. The proportion of cells exhibiting these oscillations became smaller after prolonged surface contact (Fig. 3b, Extended Data Fig. 6a), suggesting that surface sensing inhibits mNG-FimX oscillations and stabilizes polarization. To test whether mechanosensing with T4P induces polarization of the extension machinery, we visualized mNG-FimX in a *pilA^-^* mutant background, which also displayed oscillations (Fig. 3c, Supplementary Video 8). We found that the fractions of *pilA^-^* cells that showed mNG-FimX oscillations 10 and 60 min after surface contact were equal, near 90% (Fig. 3d). The distributions of oscillation frequencies between these two states were also indistinguishable (Extended Data Fig. 6b). Altogether, our results demonstrate that T4P-mediated mechanosensing at one pole locally recruits and stabilizes extension motors, thereby inducing a positive feedback onto their own activity. While several exogenous molecular compounds bias collective or single cell twitching migration^10–12^, our data shows chemical gradients are not necessary for active regulation of twitching.

PilB polarization sets the twitching direction of single cells, and PilG and PilH regulate T4P polarization to control reversals. We therefore investigated how the Chp system regulates PilB localization to control a cell’s direction of motion. We compared the mean localization profiles of PilB-mNG and mNG-FimX in WT, *pilG^-^* and *pilH^-^* backgrounds (Fig. 4a, b, Extended Data Fig. 7a). Both fusion proteins had greater polar fluorescent signal in *pilH^-^* and lower polar signal in *pilG^-^* compared to WT (Fig. 4c, d). We computed a polar localization index which measures the proportion of the signal localized at the poles relative to the total fluorescence (Extended Data Fig. 7b). About 50% of the mNG-FimX and PilB-mNG signal is found at the poles for WT, 70% of the signal is polar in *pilH^-^*, and most of it is diffuse in *pilG^-^* (Fig. 4e, g). We next computed a symmetry index that quantifies the extent of signal polarization, that is how bright a pole is compared to the other, a value of 0.5 being completely symmetric (Extended Data Fig. 7b). WT cells grown in liquid had a mNG-FimX and PilB-mNG symmetry index of about 0.6 (Fig. 4f, h). In contrast, *pilH^-^* cells were more polarized, with a symmetry index close to 0.75. Compared to WT, mNG-FimX was more symmetric in a *pilG^-^* background (Fig. 4h). We verified that the increase in expression levels in the different Chp mutants did not exacerbate PilB and FimX localization and polarization (Extended Data Fig. 8). In summary, we showed that PilG promotes polar recruitment and polarization of PilB and its regulator FimX, which is counteracted by PilH.

**Figure 4.**
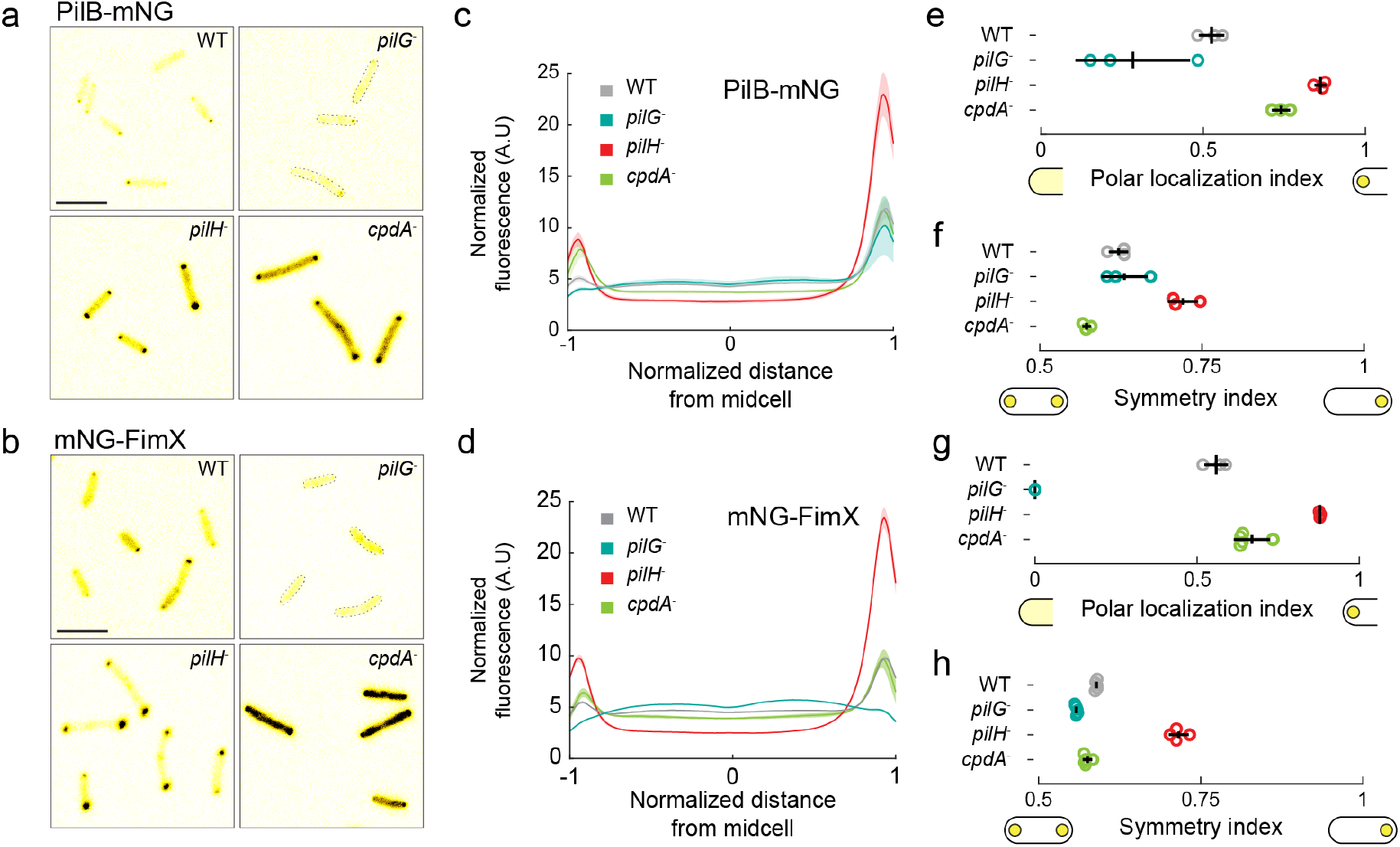
PilG and PilH control the polarization T4P extension machinery. (**a, b**) Snapshots of PilB-mNG and mNG-FimX fluorescence in WT, *pilG^-^, pilH^-^* and *cpdA^-^* background. Scale bar, 5 μm (**c, d**) Normalized fluorescence profiles along the major cell axis of the motor protein PilB and its activator FimX (Extended Data Fig.7a). Solid lines: mean normalized fluorescence profiles across biological replicates. Shaded area: standard deviation across biological replicates. (**e, g**) Polar localization index of PilB-mNG and mNG-FimX respectively, quantifying the extent of polar signal compared to a diffused configuration (Extended Data Fig.7b). An index of 0 and 1 respectively correspond to completely diffuse and polar signals. Relative to WT and *cpdA^-^*, polar localization is higher in *pilH^-^* and lower in in *pilG^-^*. (**f, h**) Symmetry index of PilB-mNG and mNG-FimX respectively, representing the ratio of the brightest pole fluorescence to the total polar fluorescence. 0.5 and 1 respectively correspond to a symmetric bipolar and a unipolar localization. *pilH^-^* has higher symmetry index than WT and *cpdA^-^*. Circles: median of each biological replicate. Black bars: (vertical) mean and (horizontal) standard deviation across biological replicates.

We then wondered how *P. aeruginosa* orchestrates two response regulators with opposing functions. Yeast and amoebae control cell polarization in response to environmental cues using spatially structured positive and negative feedback^28^. By analogy, we considered a model wherein PilG and PilH segregate to implement positive and negative feedback at distinct subcellular locations^29^. We therefore investigated the localization of functional mNG-PilG and mNG-PilH integrated at their native chromosomal loci (Fig. 5a). We found that PilG predominantly localizes to the poles (Fig. 5b, c). PilH is mainly diffuse in the cytoplasm, with only a small fraction at the poles (Fig. 5b, c). *P. aeruginosa* can therefore implement the antagonistic functions of PilG and PilH by localizing the former to the poles and the latter to the cytoplasm.

**Figure 5.**
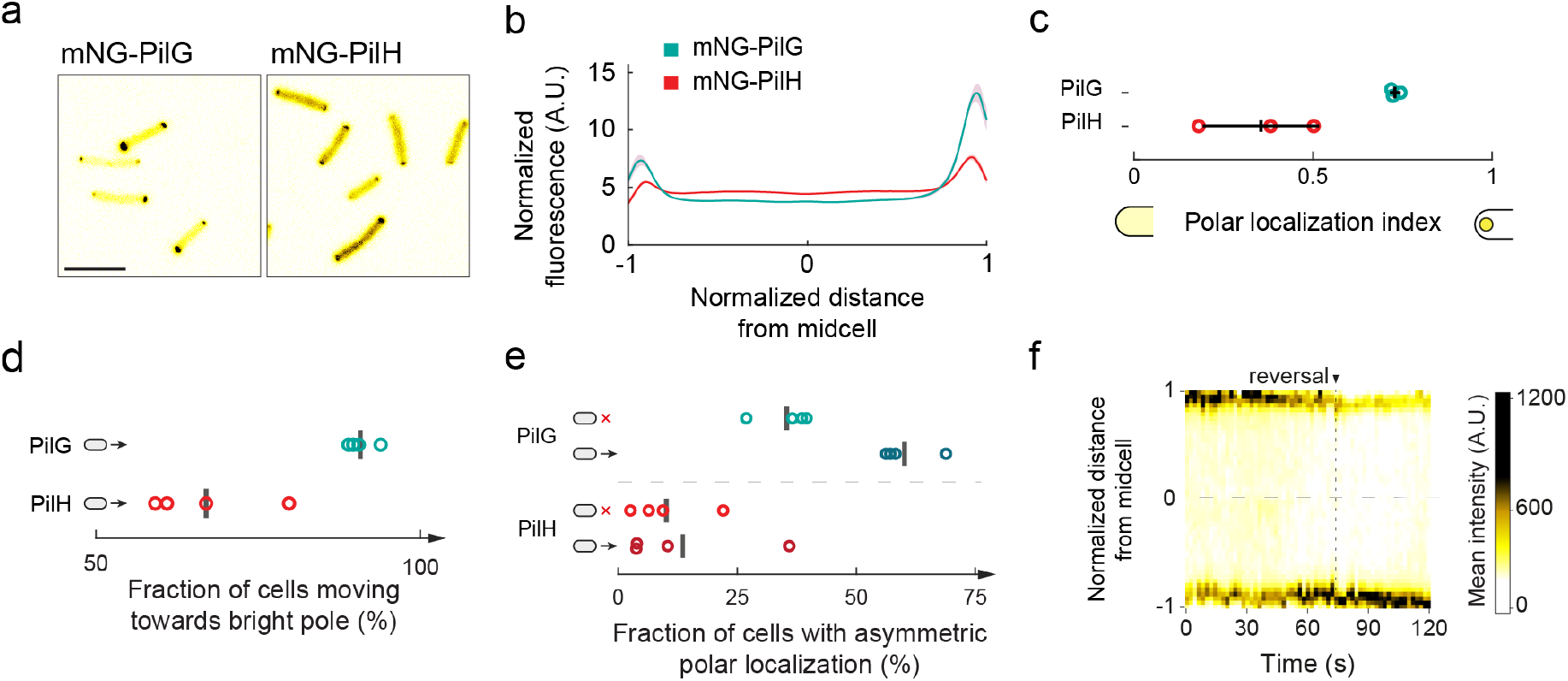
PilG and PilH dynamic localization establish a local-excitation, global-inhibition signaling landscape. (**a**) Snapshots of mNG-PilG and mNG-PilH fluorescence. Scale bar, 5 μm. (**b**) Comparison of mNG-PilG and mNG-PilH normalized mean fluorescent profiles. (**c**) The polar localization index of mNG-PilG is relatively large showing PilG is mostly polar. In contrast, mNG-PilH has a low polar localization index and is thus mostly cytoplasmic. Circles: median of each biological replicate. Black bars: (vertical) mean and (horizontal) standard deviation across biological replicates (**d**) Protein polarization relative to the twitching direction. Cells predominantly move towards the brighter mNG-PilG pole. The fraction for mNG-PilH is close to 50 %, corresponding to a random polarization relative to the direction of motion. Black bars: mean across biological replicates (**e**) Comparison of the symmetry of the polar fluorescent foci of moving cells with non-moving cells for mNG-PilG and mNG-PilH fusion proteins. There is an enrichment for mNG-PilG polar asymmetry in moving cells, but no differences in mNG-PilH. Black bars: mean across biological replicates (**f**) Kymograph of mNG-PilG in a reversing cell.

We next analyzed the relationship between a cell’s direction of migration with mNG-PilG and mNG-PilH polarization (Extended Data Fig. 9a, Supplementary Video 9). We found that 90% of cells moved towards their bright mNG-PilG pole, while only 50% did in mNG-PilH, corresponding to a random orientation relative to the direction of migration (Fig. 5d). By comparing the asymmetry of polar foci, we found that mNG-PilG signal was largely asymmetric in motile cells compared to the non-motile population (Fig. 5e). Consistent with this, in cells that reversed twitching direction, mNG-PilG localization switched to the new leading pole prior to reversal (Fig. 5f, Extended Data Fig. 9b, Supplementary Video 10). We found that the polar signal of mNG-PilH was mainly symmetric, both in moving and non-moving subpopulations (Fig. 5d). Thus, PilG, but not PilH, actively localizes to the leading pole during twitching, recapitulating the dynamic polarization of PilB and FimX. Therefore, T4P input at the leading pole activates PilG. Polar PilG drive a local positive feedback on T4P extension to maintain the direction of twitching. Cytoplasmic PilH stimulate reversals by inhibiting PilB polarization, permitting reassembly at the opposite pole. In summary, *P. aeruginosa* controls mechanotactic twitching using a local-excitation, global-inhibition signaling network architecture akin to chemotactic signaling in amoebae and neutrophils (Extended Data Fig. 10)^5,30^.

## Discussion

We discovered that *P. aeruginosa* controls the direction of twitching motility in response to mechanical signals from the motility machinery itself. This migration strategy differs from the one employed in chemotactic control of swimming motility. Chemotaxis systems control the rate at which swimming cells switch the direction of rotation of their flagella, generating successive run-and-tumbles^4,19^ or flicks^31^ that cause directional changes. However, T4P must disassemble from one pole and reassemble at the opposite in order to reverse cell movement. In essence, this tactic strategy is akin to the one of single eukaryotic cells such as amoebae and neutrophils that locally remodel their cytoskeleton to attach membrane protrusions in the direction of a stimulus^5^.

Ultimately, the ability to balance persistent forward migration with reversals optimizes *P. aeruginosa* individual twitching. Reversal may occur spontaneously or upon perturbations, for example when colliding another cell. The Chp system may also promote asymmetric piliation of upright twitching *P. aeruginosa* cells during exploratory motility^32^. We also found that the ability to reverse upon collision prevents jamming of groups of cells, supporting a model wherein the Chp system orchestrates collective migration^15^. More generally, we anticipate that other bacteria, as well as archaeal and eukaryotic species that actively migrate on surfaces leverage mechanosensation to control reversal rates and orchestrate collective motility behaviors^27^.

Beyond bacteria, eukaryotic cells also have the ability to transduce mechanical signals into cellular responses, regulating an array of physiological processes including development, immunity and touch sensation^33^. Eukaryotic cell motility is also sensitive to mechanical cues. For example, adherent mammalian cells migrate up gradients of substrate material stiffness in a process termed durotaxis^34^. We established that single cells can actively sense their mechanical environment to control motility on the timescale of seconds. Our work thus expands our view of signals activating bacterial sensing systems and more generally highlights the role of mechanics in regulating motility, be it in bacteria, archaea or eukaryotes^35^.

Altogether, the Chp system functions as a spatial sensor for mechanical input. Thus, chemotaxis-like systems can sense spatially-resolved mechanical signals, a feat that is still debated when only considering diffusible molecules as input stimuli^36^. Phototactic systems may however be an exception by conferring cyanobacteria the ability to spatially sense gradients of light^37,38^. Accordingly, the Pix phototactic system of the cyanobacterium *Synechocystis* shares signal transduction architecture with the *P. aeruginosa* Chp system by harboring two CheY-like response regulators, PixG and PixH^37,38^. There also exists a broad range of chemotaxis-like systems with even higher degrees of architectural complexity^39^. We thus highlight that their subcellular arrangement may play important functions in the mechanism by which they regulate motility.

Finally, we revealed an unexpected commonality between bacteria, and single eukaryotic cells in the way they transduce environmental signals to control polarity^5^. Amoebae and neutrophils have evolved a complex circuitry which combines positive and negative feedback loops to chemotax^28^. Positive regulators activate actin polymerization locally to drive membrane protrusion in the direction of polarization. Negative regulators inhibit actin polymerization throughout the cytoplasm to limit directional changes while also permitting adaptation^28^. Altogether, these cells establish a local activation-global inhibition landscape that balances directional persistence with adaptation^30^. By virtue of PilG and PilH compartmentalization, *P. aeruginosa* replicates the local activation-global inhibition landscape^30^. We have therefore uncovered a signal transduction architecture permitting mechanotaxis in response to spatially-resolved signals that is evolutionarily conserved across kingdoms of life.

## Acknowledgments

LT, ZAM, IV, XP, AP are supported by the SNSF Projects grant number 310030_189084. MJK is supported by the EMBO postdoctoral fellowship ALTF 495-2020. JNE, YI, RP, HM and are supported by an NIH R01 grant number AI129547 and by the Cystic Fibrosis Foundation (495008).

## Author contributions

MJK, LT, JNE and AP conceived and supervised the project. MJK, LT, YI, HM, RP, ZAM conducted the experiments. MJK, LT, IV, JN wrote the tracking code and analyzed the data. MJK, LT, JNE and AP wrote the manuscript.

