## Supplementary information for "Mechanotaxis directs *Pseudomonas aeruginosa* twitching motility"

*Corresponding authors

†These authors contributed equally


**Supplementary Information contains:**


Methods

Extended Data Figures

Extended Data Tables

Supplementary References

Supplementary video captions

### Methods

*Bacterial strains, growth conditions, and media*

All strains used in this study are listed in Extended Data Table 1. The *Pseudomonas aeruginosa* strain used was PAO1 strain ATCC 15692 (American Type Culture Collection). *Escherichia coli* strain DH5α was used for vector construction, *Escherichia coli* strain S17.1 was used for conjugation of vectors into *P. aeruginosa*. All bacterial strains were grown in LB medium (Carl Roth) at 37 °C with 290 rpm shaking. Regular solid LB media were prepared by adding 1.5 % ($wt.vol^{-1}$) agar (Fisher Bioreagents). For surface association experiments cells were plated on pre-heated solid LB media containing 1 % agarose standard (Carl Roth) and incubated at 37 °C for 3 hours. For twitching motility assays semi-solid tryptone media were prepared by autoclaving (5 $g.l^{-1}$ tryptone (Carl Roth), 2.5 $g.l^{-1}$ NaCl (Fisher Bioreagents), 0.5 % ($wt.vol^{-1}$) agarose standard (Carl Roth)). For selection of *E. coli* the following antibiotic concentrations were used: 100 µg/ml ampicillin, 10 $µg.ml^{-1}$ gentamycin. For selection of *P. aeruginosa* the following antibiotic concentrations were used: 300 $µg.ml^{-1}$ carbenicillin, 60 $µg.ml^{-1}$ gentamycin. For fluorescence visualization under correlative fluorescence and iSCAT cells were washed with PBS (BioConcept AG) and resuspended in M9 medium (VWR) containing 0.2 % of glucose.

*Strains and vector construction*

Plasmids and corresponding oligonucleotides are listed in Extended Data Table 2 and Extended Data Table 3, respectively. Gene deletions and gene fusions were constructed by two-step allelic exchange (according to [1]) using the suicide vectors pEX18Amp or pEX18Gent. In-frame deletions were generated by combining approximately 500-bp fragments of the up- and downstream regions of the designated gene, leaving only a few codons (usually around six) and verified by colony PCR and sequencing. In-frame insertions were constructed in basically the same fashion. Insertion fragments with wild-type length (e.g. for complementation) were integrated into the corresponding deletion strain. Fluorophore genes were fused to the N- or C-termini of the desired gene separated by a GGGGG linker and introduced into the wild-type chromosome. Expression and integrity of fluorescent fusions were verified by Western blotting and functionality was tested by twitching motility assays (see corresponding methods). Plasmids were constructed using standard Gibson assembly protocols[2] and introduced into *P. aeruginosa* cells by conjugative mating with *E. coli* S17.1 as donor.

*Twitching motility stab assay*

Square plastic dishes (100x100x20 mm, Sarstedt) were filled with 50 ml of 0.5 % agarose tryptone medium and dried in a flow hood for 30 min. Single colonies grown overnight on LB agar at 37 °C were stabbed with a 10 µl pipette tip through the agarose hydrogel to the bottom of the dish without much force to prevent separation of the hydrogel from the plastic. The plates were incubated at 37 °C overnight in a plastic bag. The hydrogel was then scored around the edges with a 10 µl pipette tip and removed carefully. Twitching motility was quantified by taking two measurements of the twitching zone diameters from two to eight independent stabs. The twitching diameter of wild-type or *fliC*$^{-}$ *P. aeruginosa* was used as reference.

*Western blot*

Cells were grown overnight in LB, diluted 1:100 in LB and grown until $OD_{600}$ = 1. One ml of cell suspension was harvested by centrifugation and resuspended in 100 µl of 2x loading bugger (Li-Cor), 100 mM DTT (Sigma) and at least 200 U/ml nuclease (Pierce) until the sample was no longer viscous. 10 µl of each sample was run on a sodium dodecyl sulfate-polyacrylamide gel electrophoresis (SDS-PAGE) at 200 V and transferred onto a nitrocellulose membrane (iblot 2). The membrane was incubated in PBS + 5 % milk powder for 1 h, then in 1:1000 anti-mNeonGreen antibody (Chromotek) in TBST + 5 % milk powder for 1 h, washed 3x with TBST and 1:10,000 Goat anti-Mouse IgG IRDye 800CW (Li-Cor) in TBST + 5 % milk powder for 1 h and washed 3x at room temperature. The blot was visualized with Odyssey CLx (Li-Cor).

*Phase contrast and fluorescence microscopy*

Microscopy was performed on an inverted Nikon TiE epifluorescence microscope controlled with NIS-Elements (version AR 5.02.03). Specific experimental procedures are given below. For pure phase contrast microscopy a 40x Plan APO NA 0.9 phase contrast objective was used. For fluorescence microscopy a 100x Plan APO NA 1.45 phase contrast oil objective and Semrock YFP-2427B or TxRed-A-Basic-NTE filters were used as needed. Images were acquired using a Hamamatsu ORCA Flash 4 camera. All fluorescent images were background subtracted with ImageJ (version 1.53c). Snapshot images and movies were generated with ImageJ and data were analyzed with custom scripts in Python (version 3.7.6 and 3.8.5) and MATLAB (version R2019b and R2020a), as specified in detail below.

*Correlative iSCAT fluorescence microscopy setup*

Our iSCAT microscope setup was previously described in[3]. For this study, we added a fluorescent channel as described in[4]. Briefly, we implemented a green fluorescence channel by coupling a blue laser with a wavelength of 462 nm (Lasertack, LDM-462-1400-C) to the iSCAT illumination path with a 490 nm long pass dichroic mirror (Thorlabs, DMLP490). The laser was focused on the back focal plane of the objective with a plano-convex lens (f = 500 mm, Thorlabs, LA1908-A). The fluorescent signal was then focused onto a CMOS camera (PointGrey, GS3-U3-23S6M) with a 400 mm focal length achromatic lens and a GFP filter (CWL = 525 nm, Bandwidth = 39 nm, Thorlabs, MF525-39). A shutter was placed in the illumination path of the fluorescence channel in order to prevent unnecessary illumination. The iSCAT illumination was monitored by modulating the acousto-optic deflectors' output (Extended Data Fig. 11).

*Glass coverslip and microfluidic chip preparation*

Glass coverslips (Marienfeld, 22x40 mm No 1.5) were cleaned as described in[3]. Briefly, they were washed sequentially with distilled water, ethanol, distilled water, isopropanol, distilled water, ethanol, distilled water and excess water was dried with a stream of nitrogen. For visualization, we either plasma-bonded 500 μm wide, 140 μm deep polydimethylsiloxane (PDMS) microchannels, fabricated using standard photolithography methods, or we deposited PDMS gaskets on the clean coverslips. PDMS gaskets were obtained using biopsy punches of 6 mm in diameter.

cAMP quantification using *PaQa-YFP*

The *PaQa-YFP* reporter system for cAMP measurements including a reference promoter fused to mKate2 has been previously described in detail[5]. Single colonies were grown overnight in LB-carbenicillin, diluted to $OD_{600}$ 0.05 and grown until mid-exponential phase. For surface-association growth, cells were grown for 3 h at 37 °C on LB 1 % agarose and then harvested in 1 ml LB by gentle scraping. 1 µl of culture was then loaded on a 1 % agarose-PBS pad and flipped onto a glass bottom dish (MatTek) prior to visualization. Several images in phase contrast, YFP and mKate2 channels were acquired. Images were binned 2x2 and fluorescence channels were background subtracted using a custom macro in ImageJ. Cells were segmented and the cell area and corresponding cell mean fluorescence were extracted using BacStalk[6]. Median PaQa-YFP to mKate2 fluorescent intensity ratios were computed with a custom Python script for each biological replicate. Each median was then normalized to the mean of the WT biological replicates of liquid cultures.

*iSCAT-based quantification of type IV pili number*

Single colonies were inoculated in LB, grown overnight and diluted 1:500 or 1:1000 followed by a grow period of 2 h to obtain mid exponential phase cultures. 100 µl of the cell suspension were plated on LB 1 % agarose, grown for 3 h at 37 °C and harvested in 500 µl LB by gentle scraping. Cells were diluted to $OD_{600}$ 0.02 prior to visualization. Both liquid and solid grown cells were either loaded on 500 µm x 140 µm PDMS microchannels or in 6 mm PDMS gaskets. Cells sticking to the surface were visualized without flow with iSCAT and movies were recorded at 10 fps for either 2 min, 1 min or 30 s. To reveal the interferometric component of the signal, each frame was processed as described in[3]. In some cases, we extracted the interferometric signal by differential processing. Briefly, we subtracted each frame at time t-1 to the frame at time t then performed the bandpass filtering and frame normalization as described in[3]. This technique allowed to drastically decrease strong background signal which revealed hidden floppy pili. However stationary pili disappeared in the process as they became part of the background. Individual movies were manually analyzed by counting the total number of pili in each cell. The residence time of each cell on the coverslip was also recorded. Finally, we computed a bootstrap median of the total pili number for each biological replicate. To compare strains, we then took the mean of the medians for each biological replicate within one strain as well as their standard deviation.

*Quantification of pili distributions*

To quantify pili at both cell poles, we selected flat laying cells (all strains with *fliC*$^-$ background). Pili from each pole were manually counted and recorded with a custom ImageJ macro. Probabilities were computed and plotted using Python. We focused our analysis on subset populations of cells that had either two, three or four pili and counted the total number of cells in those populations as well as their corresponding pili distribution combinations (for cells with two pili: (1|1), (2|0) (Extended Data Fig. 2a), for cells with three pili: (2|1), (3|0) (Extended Data Fig. 2b) and for cells with four pili: (2|2), (3|1) and (4|0) (Extended Data Fig. 2c)). We considered a cell to have symmetric pili distribution when they had 0 or 1 pilus difference between poles and we considered asymmetric distributions when cells had 2 and more pili difference between poles. We obtained a probability by dividing the cells in each pili distribution by the total number of cells of the corresponding population. We then compared these percentages with the percentages obtained by a random distribution (Extended Data Fig. 2d).

*Single cell twitching behavior without fluorescence*

Cells from an exponential culture grown in LB were diluted to an $OD_{600}$ of 0.2 in tryptone medium. The agarose pad was prepared by autoclaving tryptone medium 0.5 % agarose, pouring into round petri dishes and letting dry for 30 min in a flow hood. A round pad was cut out and 1 µl of the diluted cell suspension was pipetted onto the upper side of the agarose pad (i.e. the side that was not in contact with the plastic dish bottom). We note that all of the following conditions were critical in order to reproducibly allow cells to twitch as single, isolated cells: The medium composition (see Media), autoclaving the agarose medium, the initial optical density of the droplet, the volume of the droplet and pipetting on the upper side of the agarose pad. Then, without letting the droplet dry, the pad was flipped onto a microscope glass bottom dish (MatTek) and six droplets of PBS were added to the sides to prevent drying. The cells were incubated at 37 °C and imaged with phase contrast microscopy every hour at 0.2 frames per second for 5 min over 3 h.

The movies were processed with a custom ImageJ macro to ensure compatibility with the downstream analysis. If necessary, drift was also corrected using the *StackReg* plugin (version July 7, 2011;[7]) in that step. Cells were then segmented and tracked using BacStalk (version 1.8,[6]) to get the cell outline and position of the center of mass (CM) of each isolated bacterium for each frame. Subsequent analyses of the cell tracks were done with custom MATLAB scripts. For every analysis, cell tracks were categorized into moving and non-moving by using a speed threshold of 1 pixel per frame (here: 32.5 nm $s^{-1}$). The speed threshold was applied to the speed of each frame (converted from the displacement of the CM between the current and previous frame), setting the speed to 0 if below the threshold. Cells were only considered moving when the speed was unequal to zero for at least three subsequent frames over the tracked time.

In Fig. 1a, other cells in the snapshots were cut out for clarity (unmodified images: Supplementary Video 1).

<u>*Spatiotemporal cumulative displacement maps*</u>

In order to analyze twitching displacement relative to the initial cell orientation, we defined a cell orientation unit vector $\vec{t}$ from the CM to the initial leading pole (*cf.* Fig. 1a). The initial leading pole was determined by comparing the scalar products of the unit vectors from CM to poles A and B (arbitrary classification) to the normalized displacement vector $\vec{d}$ in the first frame with a speed above the speed threshold. The vectors $\vec{t}$ and $\vec{d}$ were then used to determine direction of the cell displacement $\delta$ relative to the initial leading pole, giving it a positive sign for forward (i.e. toward initial leading pole) and a negative sign for reverse (i.e. opposite to initial leading pole) movement. The spatiotemporal displacement maps (Fig. 1b) were generated by cumulating the direction-corrected displacement as a function of time, for 200 tracks that were between 31 and 61 frames long.

*Reversal frequency of isolated cells*

To count the reversal frequency of twitching cells, the scalar product between the normalized displacement vector $\vec{d}$ and the cell orientation unit vector $\vec{t}$ was determined (*cf*. Fig. 1a) and rounded for each frame (starting from the first frame with a speed above the speed threshold). This resulted in a series of numbers that correspond to movement of the cell toward the initial leading pole (1), toward the initial lagging pole (-1) or no movement (0) at that timepoint. Timepoints corresponding to no movement were removed. Cells were considered moving in the same direction (relative to the initial leading pole) as long as the sign remained the same. A reversal was counted when the sign changed, however, only if at least two subsequent frames before and after the reversal had the same sign (this was done to correct for frequent sign changes when a cell was in a non-moving phase). The reversal frequency (Fig. 1c) was calculated by dividing the sum all considered reversals by the total tracked time over all cell tracks for each biological replicate.

*Reversal frequency after collision with an obstacle (i.e. another cell)*

Because only isolated cells were tracked by BacStalk, we counted all collisions and potential subsequent reversals manually. A collision was only considered if the cell was moving for at least three frames in the same direction prior to the collision and the collision lasted for at least two frames (frame interval 5 sec). A reversal following a collision was considered only if it occurred within five frames after the collision ended. Freshly divided cells were not considered as they were never found to reverse after a collision, potentially due to the lack of functional pili basal bodies at the new pole. The frequency of reversals following a collision (Fig. 1e) was calculated by dividing the sum of all considered reversals by the sum of all considered collisions for each biological replicate.

In the snapshots of Fig. 1d (left), the moving and colliding cell was cut and placed at its original position into frame 10 s for clarity (unmodified images: Supplementary Video 3).

*Dynamic localization of fluorescent protein fusions in isolated twitching cells*

All localization studies in moving cells were performed in *fliC*⁻ background to ensure that cells are not moving by means of flagella. Cells and microscope dishes were essentially prepared as described above (*cf*. single cell twitching behavior without fluorescence). The cells were then incubated at 37 °C and imaged with fluorescence microscopy every hour at 0.2 – 0.5 frames per second for 2 min over 3 h. Cells were similarly segmented and tracked with BacStalk, and basic MATLAB analysis was performed as described above, applying a speed threshold (here: 26 - 65 nm $s^{-1}$) in order to categorize cell tracks into moving and non-moving. Next, the positions of both cell poles were determined using the cell outline obtained from BacStalk. Poles were labeled according to their position in subsequent frames. Average fluorescence intensity was measured in an area around the coordinate of each pole with a radius $r = \frac{cell\ width}{1.8}$.

Cells were then categorized according to their ratio of average fluorescence intensity between the two poles over the whole cell track ($symmetry\ ratio = \frac{intensity\ \mathrm{dim}\ pole}{intensity\ bright\ pole}$). We defined a symmetry ratio threshold (0.69) according to the distribution for mNG-PilB, which showed two subpopulations, that we considered having symmetric and asymmetric subcellular protein localization. Cells were considered symmetric with a ratio below and asymmetric with a ratio above the threshold (Fig. 2d and 5e).

In order to measure if cells moved toward the dim or the bright pole (Fig. 2e, 5d; including cells which we considered having symmetric protein localization), we defined a unit vector $\vec{e}$ from the dim to the bright pole. We then computed an alignment factor $\alpha$, which is the scalar product of the normalized displacement vector $\vec{d}$ and the fluorescence polarity vector $\vec{e}$. The alignment factor $\alpha$ represents in which direction the cell is moving relative to the bright pole. An alignment factor $\alpha > 0$ corresponds to movement toward the bright pole, with $\alpha = 1$ corresponding to movement exactly parallel to the cell length axis.

Kymographs were generated with BacStalk (version 1.8;[6]) after normalizing the cell length to 50 pixels and resampling the fluorescence intensity data with the *interp1* function in MATLAB. Because the poles were labeled inconsistently by BacStalk, the reference pole was set manually for each frame to represent the initial leading pole.

*Pili number quantification and motor proteins localization*

Overnight cultures grown in LB were diluted 1:500 to 1:1000 and grown for 3 h up to mid-exponential phase then plated for surface-association experiments as described above. Cells were harvested in 500 µl of LB by gentle scraping of the plate and loaded into PDMS gaskets. After 5 min incubation at room temperature (RT) unattached cells were washed with PBS and resuspended in M9 medium to avoid background autofluorescence of LB. Cells were visualized for 4 to 5 s with correlative iSCAT fluorescence microscopy at 100 fps in the iSCAT channel and 5 fps in the fluorescence channel. iSCAT movies were processed as described above. Further image processing and analysis was performed using custom ImageJ macros. Fluorescence images were background subtracted and iSCAT and fluorescence frames were registered using ImageJ's *Align Image by line ROI* function. The scaling factor between fluorescence and iSCAT frames was determined by the difference in effective pixel size of the two cameras. The rotation angle was computed with the two ROI lines used for registration. Pili were manually counted on iSCAT images and total fluorescence at the poles was measured on corresponding fluorescence images. Briefly, cells were selecting by the user and cell poles were automatically detected by segmenting the fluorescent cell, extracting its outline, finding its center of mass (CM) and finding the points within the outline with maximum distance from the CM. The total fluorescence at the poles was then computed by measuring the mean fluorescence within a circle at the poles with a diameter corresponding to the cell width and multiplying that mean by the area of the circle. For the statistical analysis we counted how many cells had more pili at the brightest pole and how man had more pili at the dimmest pole in each biological replicate (at least 20 cells per biological replicate). We then divided that number by the total number of cells within the biological replicate and computed the mean probability across the biological replicates for each strain. Data was plotted using Python.

*Frequency of bright pole oscillations*

Cells and microscope dishes were prepared and microscopy was performed essentially as described above (*cf.* dynamic protein localization). The only difference was that cells were grown and imaged in a heated chamber (37 °C) on the microscope at 0.2 frames per second for 3 min after 10 and 60 min. Earlier timepoints were challenging to record, because of drift and autofocus issues, that usually diminished after a few minutes after preparing the microscope slide.

To count how often the brightest fluorescence signal of a fusion protein switched from one pole to the opposite pole, a similar analysis as for the dynamic protein localization was applied to measure the average fluorescence intensity of both poles over time. Then, a series of numbers (1 or -1) was generated that corresponds to a pole (arbitrary but same over the whole track) being the bright pole (1) or the dim pole (-1) for each frame. Similar to counting reversals, a bright pole switch was counted when the sign changed. Here, no additional filters were applied. The frequency of bright pole switches (Extended Data Fig. 6) was calculated for each cell by dividing the sum of all bright pole switches by the tracked time.

*Motor protein and response regulator localization in immobilized cells*

Cells were grown, visualized and analyzed as described above (*cf*. PaQa-YFP reporter quantification). To characterize the spatial localization of the protein fusions, we segmented each cell in phase contrast and extracted their fluorescence profiles using BacStalk. Fluorescent profiles correspond to the mean pixel value of a transversal section of the cell along the mid-cell axis (Extended Data Fig. 7a). To compare cells with different expression levels, profiles were normalized by the total fluorescence of the cell (corresponding to the area under the red curve in Extended Data Fig. 7a) and rescaled by the cell length. We then computed mean profiles and standard deviations for every biological replicate. To quantify the extent of polar localization, we computed a polar localization index. To calculate this index, we integrated the fluorescence intensity at the poles defined by a circle having a diameter of a single cell width (normalized by the cell length, pink areas a and b in Extended Data Fig. 7b). We corrected this value by subtracting the fluorescence at the poles of the diffused portion of the signal corresponding to the cytoplasmic median fluorescence (hatched areas c and d in Extended Data Fig. 7b). The polar localization index was computed by dividing this value by the sum of the pole areas. A value of 0 correspond to a totally diffused protein whereas a value of 1 correspond to a totally polar protein. Similarly, we computed a symmetry index by taking the ratio between the maximum polar total fluorescence and the sum of the polar total fluorescence. A value of 0.5 corresponds to a perfectly symmetric bipolar localization whereas a value of 1 corresponds to a perfectly unipolar localization. To compare strains, we computed the mean and standard deviation across biological replicates using a custom MatLab script.

*Localization extension motors under arabinose induction*

Cells harboring the PilB-mNG expression plasmid were grown overnight in LB gentamycin. Cells were diluted 1:1000 in LB gentamycin and arabinose was added in the following concentrations: 0 %, 0.03 % and 0.1 %. The diluted cultures were grown for 3 h at 37 °C with 290 rpm shaking. Samples were then prepared, visualized and analyzed as described above.

### Data and code availability

All data are available from the corresponding authors upon request. All ImageJ, Python and MATLAB code will be uploaded to Github under the following repository: https://github.com/PersatLab/Mechanotaxis.git

### Extended Data Figures

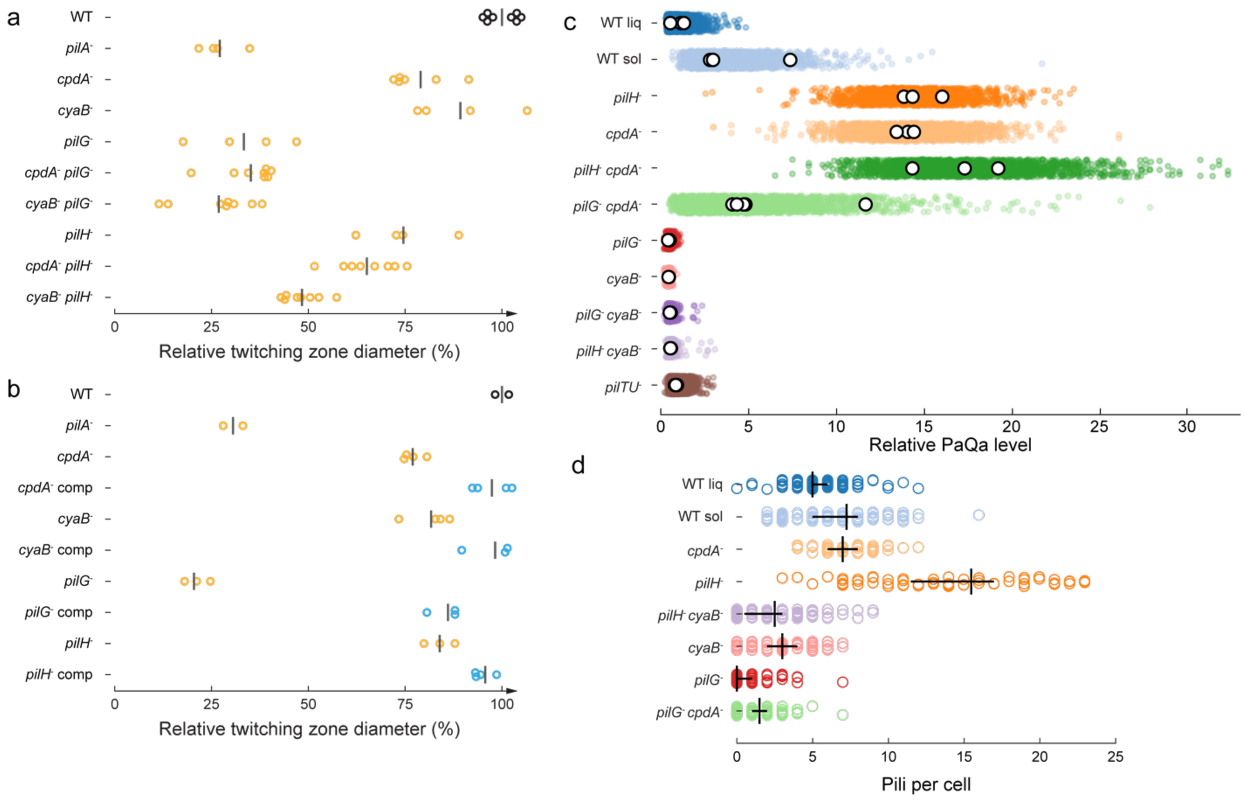


**Extended Data Figure 1: Quantification of twitching motility by the stab assay, cAMP levels and piliation of Chp mutants**. (**a**) Twitching motility as measured by the subsurface stab assay of marker-free in-frame deletions of selected Pil and Chp genes. The diameter of the zone for any given mutant is normalized by the mean twitching diameter of WT (background strain PAO1 *fliC*$^-$). Each circle corresponds to a single colony. Black bars correspond to the mean of the replicates. (**b**) Twitching zone diameters of the in-frame deletion mutants and their complemented versions. Yellow circles correspond to the in-frame deletions, blue circles correspond to the complemented mutants. Black bars correspond to the mean diameter of the replicates. Complementation of each of the in-frame deletion mutants restored the twitching zone diameter to WT levels (background strain PAO1). (**c**) cAMP levels measured by PaQa-YFP reporter fluorescence. Colored circles represent single cells, white circles correspond to median of biological replicates. All levels are normalized by the mean fluorescence intensity of the liquid-grown WT across biological replicates. All strains except WT sol are grown in liquid culture. (**d**) Quantification of T4P number in Chp and cAMP mutants by iSCAT. Each circle corresponds to the T4P number of an individual cell. Black vertical bars correspond to the boostrap mean of biological replicates, horizontal bars to their bootstrap 95 % confidence interval. All strains except WT sol are grown in liquid culture.

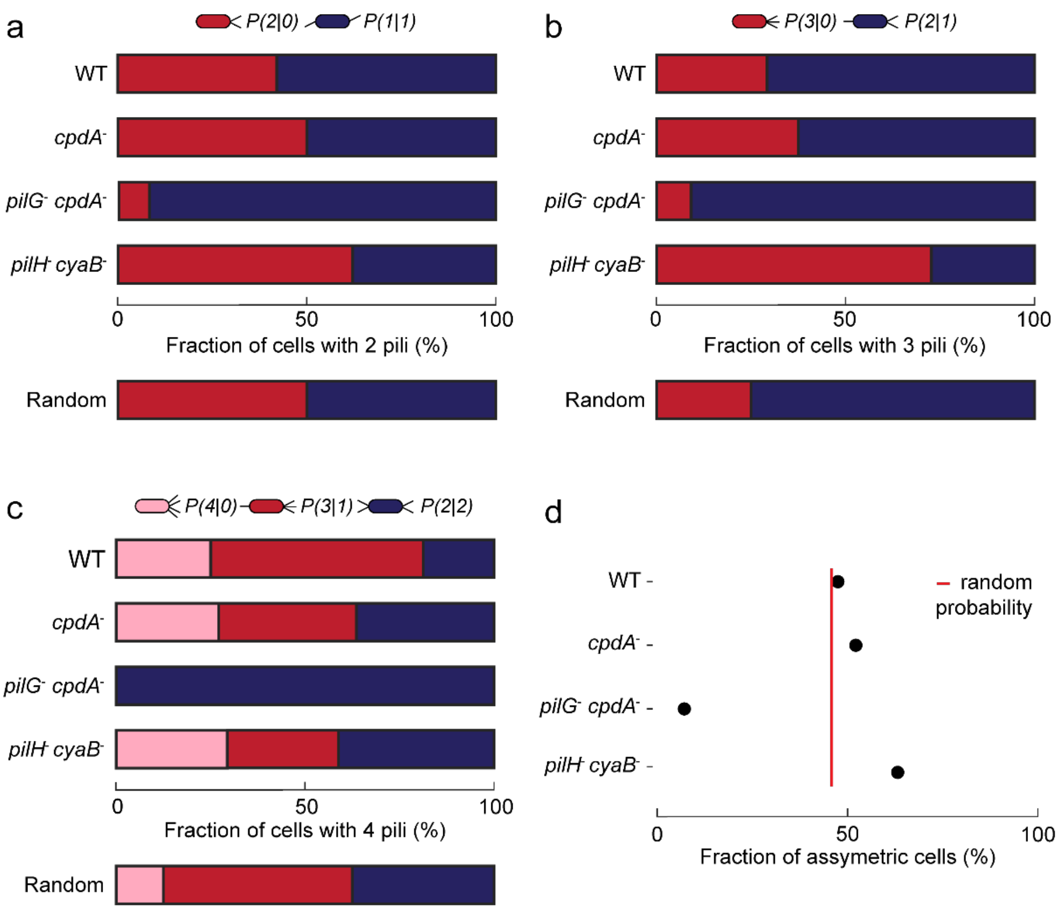


**Extended Data Figure 2: Chp controls T4P polar distribution in single cells.** We imaged single cells using iSCAT to quantify T4P at each pole. For cells that had two (**a**), three (**b**) and four (**c**) T4P, we calculated the percentage of any given T4P distribution. P(*n*/*m*) corresponds to the fraction of cells that have *n* T4P at one pole and *m* T4P at the opposite pole. For each T4P number, red corresponds to the most asymmetric distribution and blue to the most symmetric as illustrated above the graphs. As a reference, we computed the expected fractions if T4P were to be distributed randomly (shown below the mutants). WT and *cpdA*- T4P distributions are close to random. However, the distributions tend to shift toward complete symmetry in *pilG*- and complete asymmetry for *pilH*- and are independent of cAMP levels. (**d**) Cumulative fraction of the population with assymetric pili distribution in cells with two, three and four pili. *pilG*- mostly have symmetric T4P distribution whereas *pilH*- tend to polarize T4P to one pole. All cells are liquid-grown.

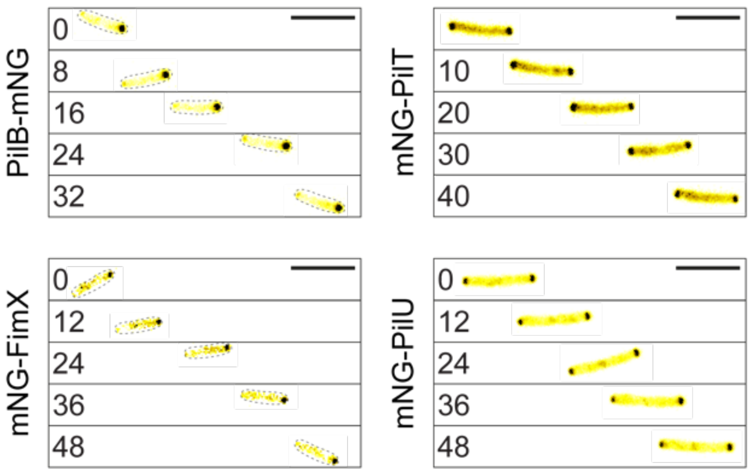


**Extended Data Figure 3**: **Timeseries of snapshots of fluorescent fusion proteins in single cells twitching forward.** PilB and FimX localize more prominently to the leading pole, while PilT and PilU tend to localize more symmerically to both poles. Images correspond to Supplementary Video 5. Scale bar, 5 μm.

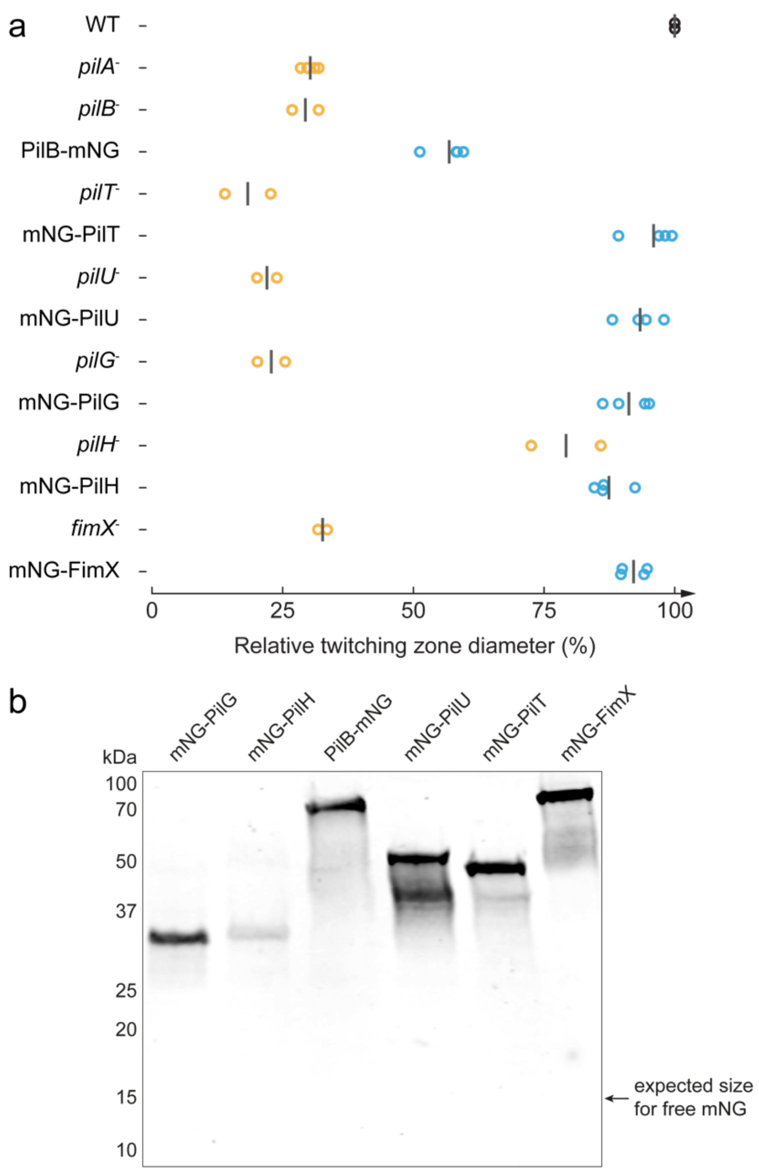


**Extended Data Figure 4**: **Functional characterization of fluorescent protein fusions to T4P motors and response regulators.** (**a**) Twitching diameters (relative to WT) as measured by the subsurface stab assay. Yellow circles correspond to in-frame deletions, blue circles correspond to chromosomally localized fluorescent fusions, black bars correspond to the mean of diameter of twitching zones. (**b**) Western blot of all mNG fluorescent fusions used in this study to verify expression levels and production of full-length protein fusion. Free mNG was not detected.

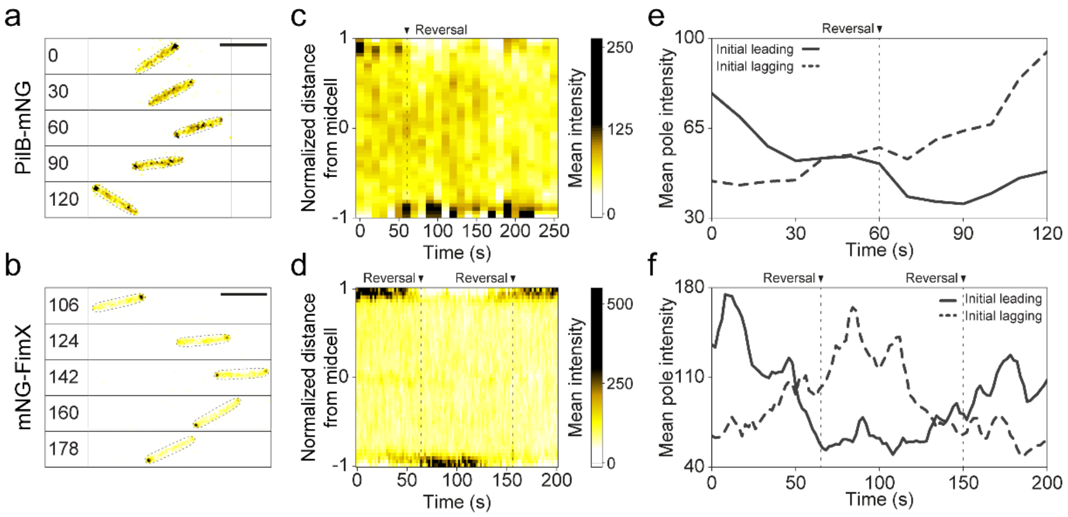


**Extended Data Figure 5: PilB-mNG and mNG-FimX switch polarization toward the new leading pole during twitching reversals.** Snapshots of fluorescence image sequences of reversing PilB-mNG **(a)** and mNG-FimX (**b**) cells (corresponding to Supplementary Video 6). (**c and d**) Corresponding kymographs during reversal showing relocalization of the fluorescent fusion proteins. (**e and f**) Fluorescent intensity traces of the leading and lagging poles during the visualization. The dynamic localization behaviour of FimX during reversals is qualitatively similar to the one described previously [8]. Scale bar, 5 μm.

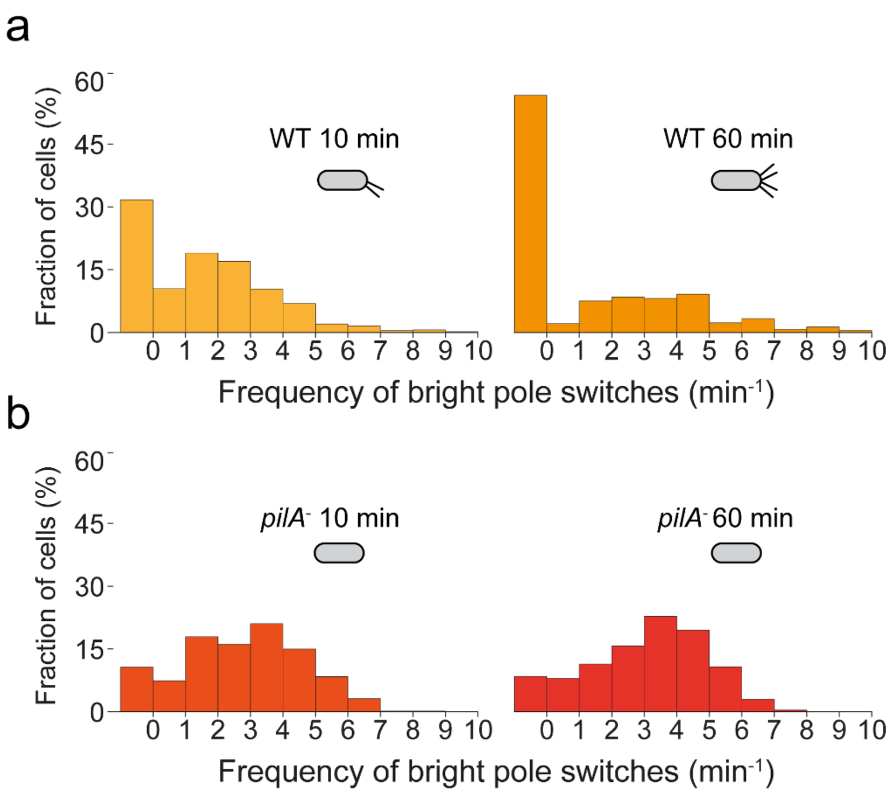


**Extended Data Figure 6: Frequency distributions of mNG-FimX oscillations upon surface contact is T4P-dependent.** (**a**) Distributions of bright foci mean switching frequency in WT 10 min (left) and 60 min (right) after surface contact. Right after surface contact, 70% of the population shows pole-to-pole mNG-FimX switches, which goes down to 50% after 1 h on the surface. (**b**) Distributions of bright focus spot switching frequency in *pilA*$^-$. Most cells show pole-to-pole switches. There is no change in the distribution of switching frequencies between cell populations that have been in contact with the surface for 10 min and 60 min.

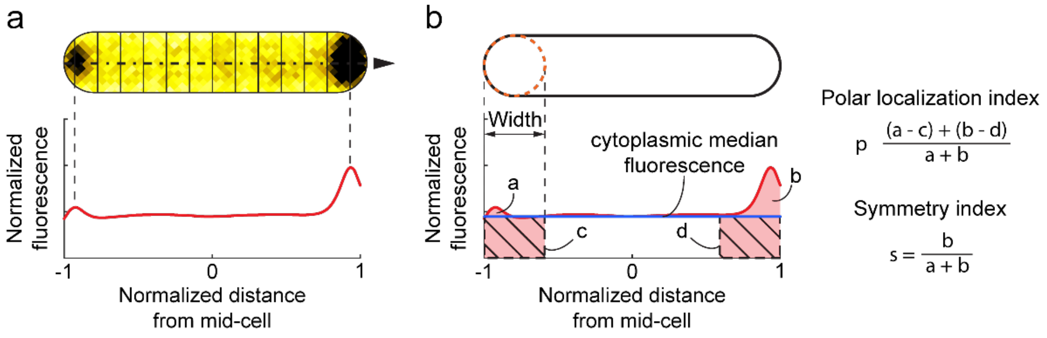


**Extended Data Figure 7**: **Computation of fluorescent profile, polar localization and symmetry incides.** (**a**) Fluorescence profiles were computed with BacStalk by taking the cross-sectional mean pixel value along the cell (dashed line). To correct the profile for differences in expression levels between cells, each was normalized by the total fluorescence intensity of the cell, represented by the area under the red curve. The cell length was also normalized so that the poles are located at -1 and 1 *x*-coordinates. (**b**) Computation of the polar localization and the symmetry indices. The pole areas (pink areas a and b) were defined by the width of the cell. The hatched areas c and d correspond to the pole areas if the protein-fusion was diffused with fluorescence levels identical to the cytoplasmic median fluorescence of the profile (blue line). To extract the polar localization index for each cell, we subtracted the diffused total fluorescence (c and d) to the actual pole total fluorescence (a and b) and divided this value by the total fluorescence at the poles (a + b). The symetry index for each cell was computed as the total fluorescence of the brightest pole b divided by the sum of the total fluorescence of the two poles a + b.

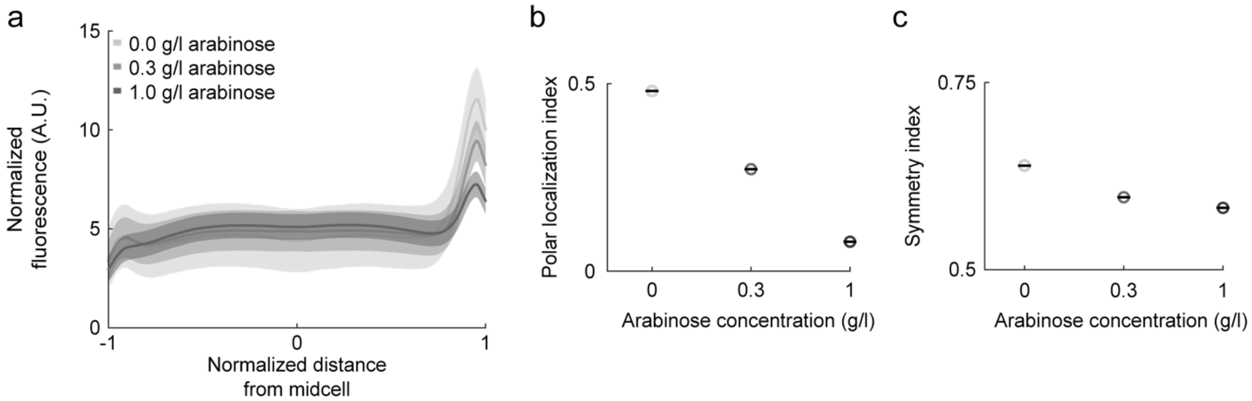


**Extended Data Figure 8: PilB-mNG polar localization and polarization as a function of expression level.** We induced the expression of PilB-mNG on a low-copy plasmid with arabinose in a *pilB*$^{-}$ background. (**a**) Mean profiles (solid lines) and standard deviation (grey areas) of a population of liquid grown cells in which PilB-mNG was induced for 3 h at three different arabinose concentrations. When normalized, the profiles flatten with increasing inducer concentration. We attribute this to the accumulation of the motor protein in the cytoplasm after T4P machineries are saturated at the poles. (**b and c**) As a result, the polar localization and symmetry indices of PilB-mNG decrease with increased concentration of inducer.

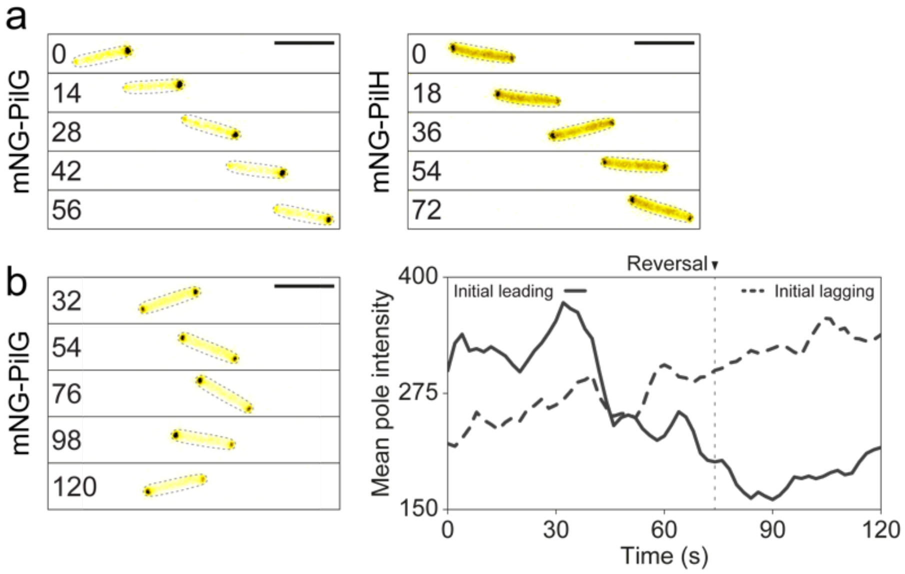


**Extended Data Figure 9: mNG-PilG and mNG-PilH localization during twitching and reversals.** (**a**) Snapshots of mNG-PilG and mNG-PilH fluorescent fusions in isolated twitching cells. PilG tends to localize more prominently to the leading pole, while PilH predominantly localizes to the cytoplasm and can be enriched at both poles without marked polarization. Snapshots taken from Supplementary Video 9. (**b**) Snapshots of a fluorescence image sequence of a reversing cell expressing mNG-PilG at its native locus (taken from Supplementary Video 10).

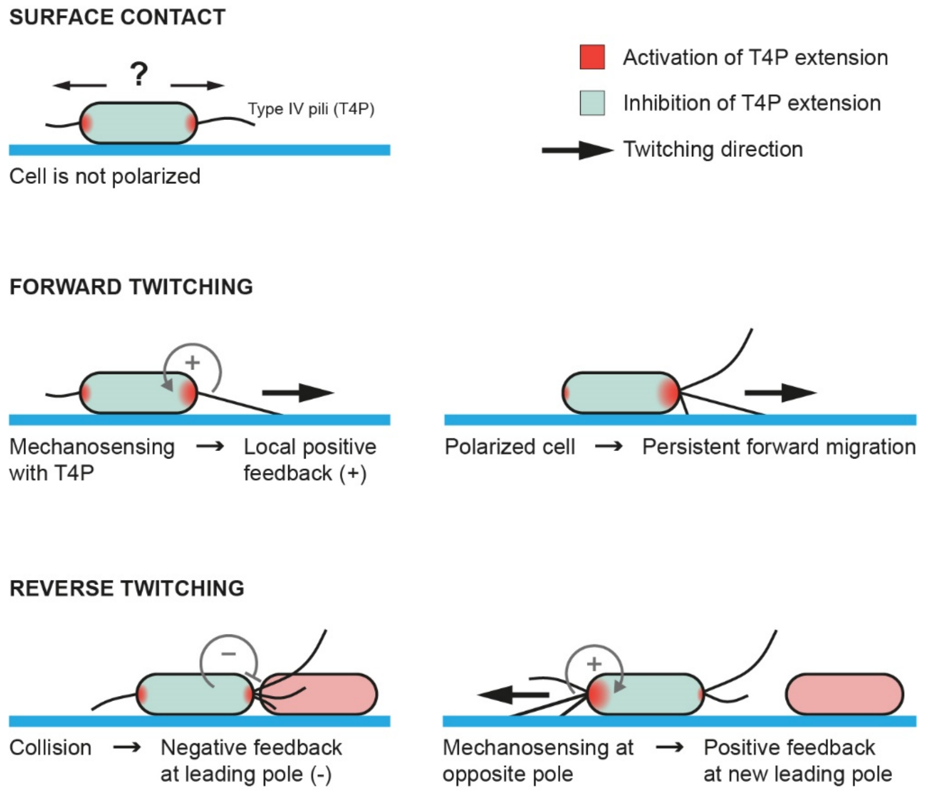


**Extended Data Figure 10: Mechanotaxis model:** After initial contact a cell explores its surrounding with random T4P distribution at both poles. Upon T4P tip attachment, Chp mechanosensing induces a positive feedback on T4P motors to favor extension at the same pole leading to polarization of the cell and persistent forward motion. After collision (or loss of T4P attachment/retraction) at the leading pole, a negative feedback down regulates T4P activity. T4P at the opposite pole can then attach and generate a positive feedback that reverses cell polarization and lead to persistent reverse twitching.

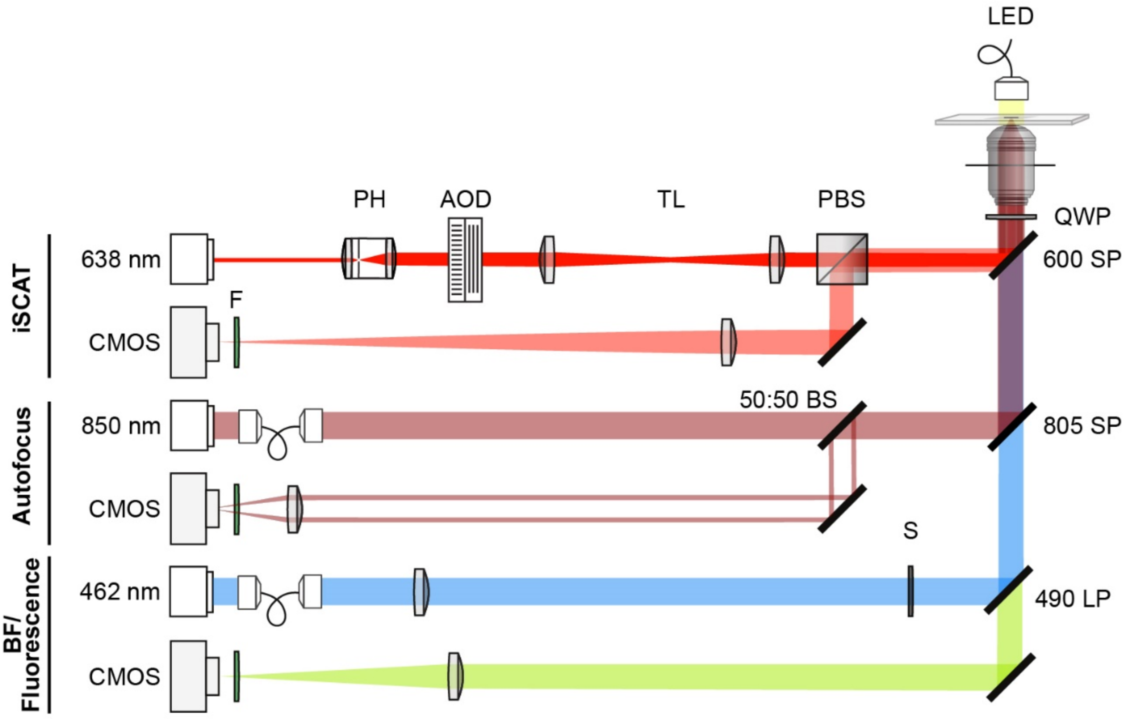


**Extended Data Figure 11: Optical layout of the correlative iSCAT fluorescence setup.** The setup is a modified version of the one described in [3] in which we added a fluorescent channel. Briefly, the fluorescence channel consists in a 462 nm blue laser spatially filtered by an optical fiber and focused to the back focal plane of the objective with a 500 mm lens and a 490 nm long pass dichroic mirror. A shutter (S) allows clipping of the beam to reduce light exposure. The fluorescent signal emitted by the sample is directed to a CMOS camera with a 400 mm imaging lens. Appropriate filters (F) were used to prevent unwanted signal reaching the detectors (iSCAT: 635 band pass (BP), Bright field/fluorescence: 525 BP, Autofocus: 850 BP).

**Extended Data Table 1:** Strains used in this study.

| **Name and relevant genotype** | **Source / Reference** | **Identifier** |
|---|---|---|
| Pseudomonas aeruginosa PAO1 | [9] | ATCC 15692 |
| Escherichia coli DH5α (hsdR rec lacZYA $\phi$80 lacZM15) | Invitrogen | Na |
| Escherichia coli strain S17.1 (thi pro hsdR recA RP4-2(Tc::Mu)(Km::Tn7)) | Stratagene | Na |
| PAO1 *pilA*$^-$ (in-frame deletion of PA4525) | [10] | 81 |
| PAO1 *pilB*$^-$ (in-frame deletion of PA4526) | [10] | 172 |
| PAO1 *pilT*$^-$ (in-frame deletion of PA0395) | [10] | 89 |
| PAO1 *pilU*$^-$ (in-frame deletion of PA0396) | [10] | 87 |
| PAO1 *pilTU*$^-$ | [10] | 171 |
| PAO1 *pilG*$^-$ (in-frame deletion of PA0408) | [10] | 226 |
| PAO1 *pilH*$^-$ (in-frame deletion of PA0409) | [11] | 178 |
| PAO1 *fimX*$^-$ (in-frame deletion of PA4959) | [12] | 885 |
| PAO1 *cpdA*$^-$ (in-frame deletion of PA4969) | [13] | 180 |
| PAO1 *cyaB*$^-$ (in-frame deletion of PA3217) | [13] | 174 |
| PAO1 *cpdA*$^-$ *pilG*$^-$ | This study | 487 |
| PAO1 *cpdA*$^-$ *pilH*$^-$ | This study | 488 |
| PAO1 *cyaB*$^-$ *pilG*$^-$ | This study | 403 |
| PAO1 *cyaB*$^-$ *pilH*$^-$ | This study | 404 |
| PAO1 *cpdA*$^-$ complementation (in-frame insertion at native locus) | This study | 1110 |
| PAO1 *cyaB*$^-$ complementation (in-frame insertion at *attB* locus) | [13] | 1091 |

| | | |
|---|---|---|
| PAO1 *pilG*⁻ complementation (in-frame insertion at native locus) | [10] | 1093 |
| PAO1 *pilH*⁻ complementation (in-frame insertion at native locus) | [10] | 1094 |
| PAO1 *fliC*⁻ (in-frame deletion of PA1092) | [10] | 177 |
| PAO1 *fliC*⁻ *cpdA*⁻ | This study | 337 |
| PAO1 *fliC*⁻ *cyaB*⁻ | This study | 326 |
| PAO1 fliC⁻ pilH⁻ | This study | 232 |
| PAO1 fliC⁻ pilG⁻ | This study | 330 |
| PAO1 fliC⁻ cpdA⁻ pilG⁻ | This study | 459 |
| PAO1 fliC⁻ cyaB⁻ pilH⁻ | This study | 436 |
| PAO1 PilB-mNG (C-terminal fluorescent fusion to mNeonGreen, GGGGG linker, native locus) | This study | 592 |
| PAO1 PilB-mNG *cpdA*⁻ | This study | 716 |
| PAO1 PilB-mNG *cyaB*⁻ *pilH*⁻ | This study | 1104 |
| PAO1 PilB-mNG *pilH*⁻ | This study | 605 |
| PAO1 PilB-mNG *pilG*⁻ | This study | 590 |
| PAO1 mNG-PilT (N-terminal fluorescent fusion to mNeonGreen, GGGGG linker, native locus) | This study | 591 |
| PAO1 mNG-PilT *pilG*⁻ | This study | 589 |
| PAO1 mNG-PilT *pilH*⁻ | This study | 648 |
| PAO1 mNG-PilT *cpdA*⁻ | This study | 773 |
| PAO1 mNG-PilU (N-terminal fluorescent fusion to mNeonGreen, GGGGG linker, native locus) | This study | 448 |
| PAO1 mNG-PilU *pilG*⁻ | This study | 604 |
| PAO1 mNG-PilU *pilH*⁻ | This study | 698 |

| | | |
|---|---|---|
| PAO1 mNG-PilU *cpdA*$^-$ | This study | 718 |
| PAO1 mNG-PilG (N-terminal fluorescent fusion to mNeonGreen, GGGGG linker, native locus) | This study | 512 |
| PAO1 mNG-PilH (N-terminal fluorescent fusion to mNeonGreen, GGGGG linker, native locus) | This study | 446 |
| PAO1 mNG-FimX (N-terminal fluorescent fusion to mNeonGreen, GGGGG linker, native locus) | This study | 421 |
| PAO1 mNG-FimX *cyaB*$^-$ *pilH*$^-$ | This study | 1102 |
| PAO1 *fliC*$^-$ PilB-mNG | This study | 316 |
| PAO1 *fliC*$^-$ PilB-mNG *cpdA*$^-$ *pilG*$^-$ | This study | 1037 |
| PAO1 *fliC*$^-$ mNG-PilT | This study | 313 |
| PAO1 *fliC*$^-$ mNG-PilU | This study | 314 |
| PAO1 *fliC*$^-$ mNG-PilG | This study | 923 |
| PAO1 *fliC*$^-$ mNG-PilH | This study | 315 |
| PAO1 *fliC*$^-$ mNG-FimX | This study | 463 |
| PAO1 *fliC*$^-$ mNG-FimX *cpdA*$^-$ | This study | 1022 |
| PAO1 *fliC*$^-$ mNG-FimX *pilH*$^-$ | This study | 965 |
| PAO1 *fliC*$^-$ mNG-FimX *pilG*$^-$ | This study | 941 |
| PAO1 fliC$^-$ mNG-FimX pilG$^-$ cpdA$^-$ | This study | 1042 |
| PAO1 *fliC*$^-$ mNG-FimX *pilA*$^-$ | This study | 1044 |
| PAO1 *pilB*$^-$ + pJN105GM-PilB-mNG (arabinose inducible) | This study | 744 |
| PAO1 *fliC*$^-$ PaQa | This study | 764 |
| PAO1 *fliC*$^-$ *pilH*$^-$ PaQa | This study | 864 |
| PAO1 *fliC*$^-$ *cpdA*$^-$ PaQa | This study | 867 |

| | | |
|---|---|---|
| PAO1 *fliC*$^{-}$ *cpdA*$^{-}$ *pilH*$^{-}$ PaQa | This study | 951 |
| PAO1 *fliC*$^{-}$ *pilG*$^{-}$ PaQa | This study | 865 |
| PAO1 *fliC*$^{-}$ *cpdA*$^{-}$ *pilG*$^{-}$ PaQa | This study | 950 |
| PAO1 *fliC*$^{-}$ *cyaB*$^{-}$ PaQa | This study | 866 |
| PAO1 *fliC*$^{-}$ *cyaB*$^{-}$ *pilH*$^{-}$ PaQa | This study | 949 |
| PAO1 *fliC*$^{-}$ *cyaB*$^{-}$ *pilG*$^{-}$ PaQa | This study | 948 |
| PAO1 *fliC*$^{-}$ *pilTU*$^{-}$ PaQa | This study | 1121 |

493 **Extended Data Table 2:** Plasmids used in this study.

| **Name and relevant information** | **Source / Reference** | **Identifier** |
|---|---|---|
| pEX100TAP (Suicide vector based on pUC19, $Amp^R$, ColE1 ori (*E. coli*), *oriT*, *sacB*, *lacZα*) | [14] | Na |
| pEX18AP (Suicide vector based on pUC18, $Amp^R$, ColE1 ori (*E. coli*), *oriT*, *sacB*, *lacZα*) | [15] | Na |
| pEX18GM (Suicide vector based on pUC18, $Gm^R$, ColE1 ori (*E. coli*), *oriT*, *sacB*, *lacZα*) | [15] | Na |
| pJN105 (arabinose-inducible plasmid based on pBBR1MCS-5, $Gm^R$, pBBR1 ori (*E. coli* and *P. aeruginosa*), *araC*-$P_{BAD}$, *lacZα*) | [16] | Na |
| pUCP18 (high-copy plasmid derived from pUC18, $Amp^R$, ColE1 ori (*E. coli*), R1822-based ori (*P. aeruginosa*)) | [17] | Na |
| pEx100TAP-*cpdA*$^-$ (Suicide vector for marker-free in-frame deletion of PA4969) | [13] | pJTW033 |
| pEX18GM-*cpdA*$^-$ (Suicide vector for marker-free in-frame deletion of PA4969) | This study | pMK019 |
| pEx100TAP-*cyaB*$^-$ (Suicide vector for marker-free in-frame deletion of PA3217) | [13] | pJTW031 |
| pEX18GM-*cyaB*$^-$ (Suicide vector for marker-free in-frame deletion of PA3217) | This study | pMK018 |
| pEx100TAP-*pilG*$^-$ (Suicide vector for marker-free in-frame deletion of PA0408) | [10] | PJB118 |
| pEx100TAP-*pilH*$^-$ (Suicide vector for marker-free in-frame deletion of PA0409) | [10] | PJB119 |
| pEx100TAP-*fliC*$^-$ (Suicide vector for marker-free in-frame deletion of PA1092) | [10] | pJB215 |

| | | |
|---|---|---|
| pEx100TAP-PilB-mNG (Suicide vector for marker-free in-frame insertion of *p*ilB C-terminus fused with mNeonGreen, separated by a GGGGG linker) | This study | pXP121 |
| pEx100TAP-mNG-PilT (Suicide vector for marker-free in-frame insertion of *p*ilT N-terminus fused with mNeonGreen, separated by a GGGGG linker) | This study | pXP118 |
| pEx100TAP-mNG-PilU (Suicide vector for marker-free in-frame insertion of *p*ilU N-terminus fused with mNeonGreen, separated by a GGGGG linker) | This study | pXP119 |
| pEx100TAP-mNG-PilH (Suicide vector for marker-free in-frame insertion of *p*ilH N-terminus fused with mNeonGreen, separated by a GGGGG linker) | This study | pXP125 |
| pEx100TAP-mNG-FimX (Suicide vector for marker-free in-frame insertion of *fimX* N-terminus fused with mNeonGreen, separated by a GGGGG linker) | This study | pXP186 |
| pJN105GM-PilB-mNG (arabinose-inducible *p*ilB C-terminally fused with mNeonGreen, separated by a GGGGG linker) | This study | pXP377 |
| pUCP18-PaQa (fluorescent reporter for cAMP level: YFP controlled by *PaQa* promoter (PA1867 and PA1868) and mKate2 controlled by *rpoD* promoter (PA0576) as reference. | [5] | pAP02.2 |

**Extended Data Table 3:** Oligonucleotides used in this study.

| Name and relevant information | Source / Reference | Identifier |
|---|---|---|
| EcoR1_cyaB_F (cloning pMK018; GCT ATG ACC ATG ATT ACG GCC GGT GTT CCT GTA TGT CG) | This study | oMK054 |
| OL_cyaB_KO_R (cloning pMK018; CAT GAA GCC TGT CAT CCT CTA AGT TCG TCG AAC G) | This study | oMK055 |
| OL_cyaB_KO_F (cloning pMK018; AGA GGA TGA CAG GCT TCA TGC GCT GGA GAG G) | This study | oMK056 |
| Xba1_cyaB_R (cloning pMK018; CAT GCC TGC AGG TCG ACT TCG CCG AGT TCT ACC CCT ACT ACC) | This study | oMK057 |
| seq fwd del cyaB (check pMK018; TGA TGA CGA GCG TTT CCA CAG T) | This study | oLT021 |
| seq rev del cyaB (check pMK018; GCG TCG ATA CCG AAC TGT TCC AT) | This study | oLT022 |
| XbaI_cpdA_F (cloning pMK019; CAT GCC TGC AGG TCG ACT GAC CTG CGA AGC GAA CTA CG) | This study | oMK058 |
| OL_cpdA_KO_R (cloning pMK019; GTA TCC GGC TGA CAA GGG GCC GTC TCC) | This study | oMK059 |
| OL_cpdA_KO_F (cloning pMK019; CCC TTG TCA GCC GGA TAC TGA CAT GCC CC) | This study | oMK060 |
| EcoRI_cpdA_R (cloning pMK019; GCT ATG ACC ATG ATT ACG CCT GGT AGC GTT CGG GAC G) | This study | oMK061 |

| | | |
|---|---|---|
| seq fwd cpda KO (check pMK019; GTC TGG CCG TTG GAA GAT G) | This study | oLT009 |
| seq rev cpda KO (check pMK019; AAA AGT CGG TGT CGC GC) | This study | oLT010 |
| PilB region in PAO1 FWD (1) (cloning pXP121; ACC CTG TTA TCC CTA ACT AGC CTG TGG GGC GAG AAG ATC G) | This study | oXP175 |
| PilB region in PAO1 FWD (2) (cloning pXP121; GTA TGG ATG AAT TGT ATA AAT AAT CCA TGG CGG ACA AAG C) | This study | oXP176 |
| PilB region in PAO1 REV (1) (cloning pXP121; X GAG ACT CCG CCT CCG CCT CCA TCC TTG GTC ACG CGG TTG A) | This study | oXP177 |
| PilB region in PAO1 REV (2) (cloning pXP121; GGA TAA CAG GGT AAT ACT AGT GTA GAC GAT GCC TCC GAA A) | This study | oXP178 |
| G5linker mNeonGreen FWD (cloning pXP121; TCA ACC GCG TGA CCA AGG ATG GAG GCG GAG GCG GAG TCT C) | This study | oXP179 |
| G5linker mNeonGreen REV (cloning pXP121; GCT TTG TCC GCC ATG GAT TAT TTA TAC AAT TCA TCC ATA CCC ATT ACA TCA GTA AAA GC) | This study | oXP180 |
| Seq PilB_mNeonGreen fw (check pXP121; ACG GAG TCG AGC GCA TCG) | This study | oXP226 |
| Seq PilB_mNeonGreen rev (check pXP121; GCG CAA CCT GGC CAA GTA C) | This study | oXP227 |
| PilT PilU region in PAO1 FWD (1) (cloning pXP118; ACC CTG TTA TCC CTA ACT AGT TCG CGC AGG CGG GCG AAC G) | This study | oXP157 |

| | | |
|---|---|---|
| PilT PilU region in PAO1 FWD (2) (cloning pXP118; ATA AAG GAG GCG GAG GCG GAG ATA TTA CCG AGC TGC TCG CCT T) | This study | oXP158 |
| PilT PilU region in PAO1 REV (1) (cloning pXP118; TCT TCT TCA CCT TTA GAG ACC ATG GGA CTC CCC AAT TAC AAG CAA GCA) | This study | oXP159 |
| PilT PilU region in PAO1 REV (2) (cloning pXP118; GGA TAA CAG GGT AAT ACT AGG TCT TCG CCG CCG AGG TGG T) | This study | oXP160 |
| G5linker mNeonGreen FWD (cloning pXP118; TGT AAT TGG GGA GTC CCA TGG TCT CTA AAG GTG AAG AAG ATA ATA TGG C) | This study | oXP161 |
| G5linker mNeonGreen REV (cloning pXP118; GCG AGC AGC TCG GTA ATA TCT CCG CCT CCG CCT CCT TTA T) | This study | oXP162 |
| Seq mNeonGreen_PilT fw (check pXP118; GAC TGC GAA GGC GCC TTG G) | This study | oXP220 |
| Seq mNeonGreen_PilT rev (check pXP118; CTT CGC CGA TGG CTG CCT C) | This study | oXP221 |
| PilT PilU region in PAO1 FWD (1) (cloning pXP119; ACC CTG TTA TCC CTA ACT AGA CGA ATC GAA GAA GTG CCT G) | This study | oXP163 |
| PilT PilU region in PAO1 FWD (2) (cloning pXP119; ATA AAG GAG GCG GAG GCG GAG AAT TCG AAA AGC TGC TGC GCC) | This study | oXP164 |
| PilT PilU region in PAO1 REV (1) (cloning pXP119; TCT TCT TCA CCT TTA GAG ACC ATG ATG TTC TCG CTC ACT C) | This study | oXP165 |

| | | |
|---|---|---|
| PilT PilU region in PAO1 REV (2) (cloning pXP119; GGA TAA CAG GGT AAT ACT AGG CAC CAG TTG CTG GGC GAC G) | This study | oXP166 |
| G5linker mNeonGreen FWD (cloning pXP119; TGT GAG TGA GCG AGA ACA TCA TGG TCT CTA AAG GTG AAG AAG ATA ATA TGG C) | This study | oXP167 |
| G5linker mNeonGreen REV (cloning pXP119; GCG CGC AGC AGC TTT TCG AAT TCT CCG CCT CCG CCT CCT TTA T) | This study | oXP168 |
| Seq mNeonGreen_PilU fw (check pXP119; GCG GCG ATG CTC GAT TAC CTG) | This study | oXP222 |
| Seq mNeonGreen_PilU rev (check pXP119; CCC TTG CGG ATC AGG TCG G) | This study | oXP223 |
| PilG PilH ChpA region in PAO1 FWD (2) (cloning pXP125; ATA AAG GAG GCG GAG GCG GAG CTC GTA TTT TGA TTG TTG ATG ACT CTC CGA CC) | This study | oXP232 |
| PilG PilH ChpA region in PAO1 REV (1) (cloning pXP125; TCT TCT TCA CCT TTA GAG ACC ATG GGA TCC CCA TCA CGA A) | This study | oXP232 |
| mNeonGreen G5 linker FWD (cloning pXP125; TTC GTG ATG GGG ATC CCA TGG TCT CTA AAG GTG AAG AAG ATA ATA TGG C) | This study | oXP234 |
| mNeonGreen G5 linker REV (cloning pXP125; TCA ACA ATC AAA ATA CGA GCT CCG CCT CCG CCT CCT TTA T) | This study | oXP235 |
| PilG PilH ChpA region in PAO1 FWD (2) (cloning pXP125; ACA AGG GAG GCG GAG | This study | oXP236 |

| | | |
|---|---|---|
| GCG GAG CTC GTA TTT TGA TTG TTG ATG ACT CTC CGA CC) | | |
| PilG PilH ChpA region in PAO1 REV (1) (cloning pXP125; AGC TCC TCG CCC TTG CTC ACC ATG GGA TCC CCA TCA CGA A) | This study | oXP237 |
| Sq pilG ko fwd (check pXP125; CGT GGC GAA GAA CTT CTC G) | This study | oLT042 |
| check_pilGH (check pXP125; TTT ATA CGG CGA CGT CGA GG) | This study | oMK052 |
| FimX region in PAO1 FWD (1) (cloning pXP186; TAT GAC CAT GAT TAC GAA TTC TGC CCA TCG TCA ACC AC) | This study | oXP438 |
| FimX region in PAO1 REV (1) (cloning pXP186; TCT TCA CCT TTA GAG ACC ATG GAA AGG GCT CAG TCC) | This study | oXP439 |
| mNeonGreen G5 linker FWD (cloning pXP186; ACT GAG CCC TTT CCA TGG TCT CTA AAG GTG AAG AAG ATA ATA T) | This study | oXP440 |
| mNeonGreen G5 linker REV (cloning pXP186; CCG CCG CCC GAG CCC CCG CCG CCT TTA TAC AAT TCA TCC ATA CCC ATT A) | This study | oXP441 |
| FimX region in PAO1 FWD (2) (cloning pXP186; CGG GGG CTC GGG CGG CGG GGG CTC GAT GGC CAT CGA AAA GAA AAC CAT C) | This study | oXP442 |
| FimX region in PAO1 REV (2) (cloning pXP186; TGC CTG CAG GTC GAC TCT AGA GGC GCC CTG GTC GCA) | This study | oXP443 |
| seq FimX N' fusion fw (check pXP186; AGG CCC ACG CAC AGG) | This study | oXP456 |

| | | |
|---|---|---|
| seq FimX N' fusion rev (check pXP186; GGT GCA AAC CAG TTC GG) | This study | oXP457 |
| EcoRI pilB fwd (cloning pXP377; AAA GAA TTC ATG AAC GAC AGC ATC CAA CTG) | This study | oXP909 |
| mNG in pJN105 (rev) (cloning pXP377; CCG CTC TAG ATC ATT TAT ACA ATT CAT CCA TAC) | This study | oXP616 |

**Supplementary video captions**

**Supplementary video 1:** Phase contrast image sequences of *P. aeruginosa* cells twitching forward and reversing. Videos corresponding to Fig. 1 a. Scalebar, 2 μm.

**Supplementary video 2:** Phase contrast image sequence showing *P. aeruginosa cpdA*$^-$ *pilG*$^-$ cells moving back and forth, without much net migration. Scalebar, 10 μm.

**Supplementary video 3:** Phase contrast image sequence of reversal behaviors upon collisions. A WT *P. aeruginosa* cell collides with another cell, then reverses. A *pilH*$^-$ cell collides with another cell but doesn't reverse. Videos correspond to images in Fig. 1d. Scalebar, 2 μm.

**Supplementary video 4:** Twitching *P. aeruginosa* WT, *pilH*$^-$ and *cpdA*$^-$ *pilG*$^-$ cells at high cell density Phase contrast image sequence corresponding to Fig. 1f. Scalebar, 50 μm.

**Supplementary video 5:** Fluorescent localization of PilB-mNG, mNG-FimX, mNG-PilT and mNG-PilU in twitching cells. PilB-mNG, mNG-FimX localizes more prominently to the leading pole. Fluorescence microscopy image sequence corresponding to Extended Data Fig. 3. The arrow in the first frame indicates the cell depicted in the corresponding figure. Playback speed, 20x. Scalebar, 10 μm.

**Supplementary video 6:** Fluorescence microscopy image sequence of PilB-mNG and mNG-FimX during reversal. Videos correspond to Extended Data Fig. 5. Playback speed, 20x. Scalebar, 2 μm.

**Supplementary video 7:** Fluorescence microscopy image sequence corresponding to Fig. 2a. mNG-FimX oscillates between poles in a non-moving cell 10 min after touching the surface. Playback speed, 50x. Scalebar, 2 μm.

**Supplementary video 8:** Fluorescence microscopy image sequence corresponding to Fig. 2c. In *pilA*$^-$, mNG-FimX keeps oscillating between poles in a non-moving cell even 60 min after touching the surface. Playback speed, 50x. Scalebar, 2 µm.

**Supplementary video 9:** Fluorescent localization of mNG-PilG and mNG-PilH in twitching cells. Image sequence corresponding to Extended Data Fig. 9a. mNG-PilG localizes more prominently to the leading pole. The arrow in the first frame indicates the cell depicted in the corresponding figure. Playback speed, 20x. Scalebar, 10 µm.

**Supplementary video 10:** mNG-PilG fluorescence during a reversal. mNG-PilG localizes to the new leading pole. Fluorescence microscopy image sequence corresponding to Extended Data Fig. 9b. Playback speed, 20x. Scalebar, 5 µm.
